# Cannibalism Shapes Biofilm Structure and Composition in *Bacillus subtilis*

**DOI:** 10.1101/2025.03.21.644447

**Authors:** Lena Friebel, Jan-Philipp Knepper, Nathalie Sofie Becker, Gorkhmaz Abbaszade, Kathrin Stückrath, Jens Soltwisch, Susann Müller, Klaus Dreisewerd, Thorsten Mascher

## Abstract

In *Bacillus subtilis* colony biofilms, phenotypic diversification confers tissue-like properties and enhanced competitive fitness within a structural framework that allows both colony expansion and long-term survival via endospore formation. Cannibalism is thought to delay sporulation by enabling one subpopulation to produce the sporulation delay protein SDP, the sporulation killing factor SKF and the epipeptide EPE. These toxins are thought to lyse susceptible nonproducers, thereby releasing nutrients to prevent premature sporulation. However, the molecular mechanisms orchestrating this bacterial programmed cell death during biofilm development are poorly understood. Here, we comprehensively characterized mutants defective in either toxin production or the corresponding autoimmunity by a multiscale approach, combining luminescence reporters, colony biopsy, multi-parameter flow cytometry and MALDI-mass spectrometry imaging to resolve cannibalism function and distribution. The toxins are produced in distinct, only partially overlapping areas of the colony and interdepend in their spatial distribution. Both EPE and SDP, but not SKF, are crucial for delaying sporulation. Loss of EPE or SDP autoimmunity resulted in severe morphological changes and stress-induced occurrence of suppressor mutants. The absence of all three toxins led to small, hyper-sporulating colonies with excessive wrinkle formation, indicating that cannibalism is essential for maintaining biofilm structure and lateral expansion. Our results provide the first evidence for the complex interactions between the cannibalism toxins that shape biofilm architecture through bacterial programmed cell death. Localized toxin production and their spatial distribution affect the spatiotemporal organization, morphology and subpopulation dynamics within *B. subtilis* biofilms.

**Importance:** Programmed cell death (PCD) is a ubiquitous and crucial mechanism to structure eukaryotic multicellular tissues. PCD-like processes have also been described in bacteria, but their contribution to the multicellular development is poorly understood. Cannibalism in *Bacillus subtilis* has been described as a sporulation delay strategy, in which one subpopulation produces antimicrobial peptides that kill susceptible nonproducing siblings. Their lysis is thought to release nutrients that delay the sporulation in the producing subpopulation. This study comprehensively analyses the role of the three cannibalism toxins in shaping colony biofilms. By combining MALDI-mass spectrometry imaging, colony biopsy, flow cytometry, and luminescence reporters, we demonstrate that cannibalism toxins are crucial for biofilm structure. They show a discrete and interdependent localization within the biofilm. While cannibalism inhibits sporulation and causes severe envelope stress within colonies, our data challenges the established role of cannibalism-dependent killing as the mechanism behind this sporulation delay.

## Introduction

Biofilms, a collective mode of bacterial life, appeared over three billion years ago on our planet ^1^. The ability to form these structured, multicellular aggregates likely evolved as a survival strategy in response to the harsh environmental conditions of early Earth. It allowed bacterial cells to thrive and survive in a self-produced, matrix-protected, and surface-adherent microenvironment in which collective behaviors and differentiated multicellular traits could develop ^2^.

The soil-dwelling, Gram-positive bacterium *Bacillus subtilis* is a model organism for studying (multi)cellular differentiation and development in complex biofilms. It can remarkably transition from a motile to a sessile lifestyle, forming dormant endospores within the context of differentiated biofilms ^3^. *B. subtilis* can form different biofilms, depending on the environment: pellicle biofilms at the liquid-air interface, colony biofilms at the solid-air interface, and submerged surface-attached biofilms ^4^. Within *B. subtilis* biofilms, different cell types emerge in a spatiotemporally controlled manner during differentiation in response to specific microenvironments, e.g., based on the availability of nutrients ^5^. These processes are guided by a tightly regulated gene expression program, which is influenced by extracellular cues, stochastic gene expression, and genetic noise ^6–9^. A key player in genetic regulation is the master regulator of sporulation, Spo0A, which orchestrates cell differentiation once external stimuli propose harsh living conditions that favor the formation of endospores ^10–13^. Spo0A becomes activated through a multicomponent phosphorelay, resulting in increasing levels of Spo0A∼P over time ^14,15^. Consequently, it gradually silences regulatory repression of its counterplayer, the transition state regulator AbrB, which suppresses premature activation of stationary phase traits: increasing cellular Spo0A∼P levels represses the expression of *abrB* ^16^. Low levels of Spo0A∼P trigger the differentiation of cells into producing the extracellular matrix (ECM), which is essential for biofilm formation ^17,18^. This subpopulation then displays elevated levels of matrix-associated genes in a SinR-dependent manner ^19^. The extracellular matrix comprises exopolysaccharides, amyloid TasA fibers, and the hydrophobin-like protein BslA, which forms a hydrophobic biofilm coating ^4^. Interestingly, matrix-producing cells also exhibit characteristics of cannibalistic cells ^20^.

Cannibalism is a social behavior in *B. subtilis* that occurs in the early sporulation phase at low Spo0A∼P levels ^21^. A subset of cells is thought to produce cannibalism toxins that will lyse neighboring non-cannibalistic cells to release nutrients, which may help delay commitment to sporulation in the cannibalistic cells. Cannibalism is determined by expressing the operons encoding the sporulation killing factor SKF, the sporulation delaying protein SDP, and the more recently discovered epipeptide EPE ^17,22,23^.

The *skfABCEFGH* operon contains the structural gene for the SKF toxin (*skfA*), genes for the toxin maturation machinery (*skfB*, *skfC*), an ATP-binding cassette (ABC) transporter that mediates autoimmunity to SKF (*skfEF*), as well as two genes of yet unknown function (*skfG*, *skfH*) ^23,24^. The *sdpABC*-*sdpIR* locus consists of the toxin-encoding gene *sdpC*; the gene products of *sdpA* and *sdpB* are required for toxin maturation, while *sdpIR* encodes the autoimmunity, with SdpR being a transcriptional repressor and SdpI an SDP-sequestering lipoprotein ^25^. Within the *epeXEPAB* operon, *epeX* encodes the prepropeptide for a secreted antimicrobial peptide (EPE), *epeE* encodes a radical S-adenosyl-L-methionine (SAM) epimerase, which is necessary for maturation and processing of the EPE toxin, together with the transmembrane peptidase, encoded by *epeP*. The genes *epeAB* encode an ABC transporter that mediates autoimmunity against intrinsically produced EPE ^22^. These toxins, classified as ribosomally synthesized and post-translationally modified peptides (RiPPs), disrupt cell membrane integrity. While the exact mode of action of SKF remains unclear, SDP and EPE have been extensively studied. SDP collapses the proton motive force and triggers autolysis ^26^, while EPE dissipates membrane potential, reduces membrane fluidity, and induces lipid domain formation ^22^. In response to these destructive molecules, *B. subtilis* mounts a defense strategy known as the cell envelope stress response (CESR) ^27,28^. EPE primarily activates the two-component system LiaRS, which upregulates the *liaIH* operon to confer resistance against extrinsically applied EPE ^29^. BceRS and PsdRS respond to SDP and primarily SKF ^24^. Both two-component systems regulate ABC transporters, BceAB and PsdAB, respectively, which provide resistance against antimicrobial peptides ^30^.

Cannibalism is a bacterial form of programmed cell death (PCD) ^31^. It is pivotal in structuring bacterial colonies by facilitating wrinkle formation ^32^. These wrinkles result from vertical buckling movements driven by localized cell death, alleviating the mechanical stress associated with high cell densities and rapid cell division. Wrinkle formation depends on surface adhesion and extracellular matrix production, as matrix-deficient mutants fail to develop these structural features ^33^. These interconnected wrinkles enhance biofilm function by creating nutrient- and waste-transporting channels and improving overall biofilm physiology ^34,35^. While the biophysical processes of biofilm formation are increasingly understood, very little is known about the relationship between structural organization and differentiation processes, such as cannibalism, on a molecular and cellular level - this knowledge gap results from the technological challenges in studying bacterial multicellular aggregates due to size restrictions. Understanding the interdependence of the multicellular ‘form’ and the emerging ‘functions’ of bacterial biofilms requires resolving macroscopic structures, such as colony biofilms, at (near to) single-cell resolution. Recent advancements in instrumentation and imaging technology allow the implementation of multiscale, high-resolution approaches to study differentiated microbial communities ^36^. Matrix-assisted laser desorption ionization mass spectrometry imaging (MALDI-MSI) has seen key improvements concerning analytical sensitivity and spatial resolution in recent years ^37–42^, which now allow culturing colony biofilms on a suitable mixed cellulose ester membrane and analysis of their chemical composition at high resolution. Another powerful technique for examining bacterial colonies at single-cell level and investigating phenotypic heterogeneity is flow cytometry. Biopsies can be collected from different regions within colony biofilms and then analyzed using flow cytometry to explore spatiotemporal changes in phenotypic traits. This biopsy approach has revealed considerable heterogeneity among cells within bacterial colonies, highlighting variations in cell cycle phases, the ratio of live to dead cells, and the prevalence of different spore types ^43^. These approaches allow for insights into the complex structural organization of biofilms and help identify subpopulations and the function of individual cells within such structures and subpopulations ^36^.

In this study, we investigated the role of cannibalism in the spatial and temporal organization of *B. subtilis* colony biofilms. By combining multiparameter flow cytometry and imaging mass spectrometry with genetic alterations, we analyzed the distribution and dynamics of cannibalism toxins and the resulting distribution of different cell types within biofilms. This approach allowed us to elucidate how cannibalism-induced PCD contributes to biofilm architecture and the resulting emerging multicellular function.

## Results

### Cannibalism affects the morphology of *B. subtilis* colony biofilms

Cannibalism, a differentiation strategy that involves sacrificing one subpopulation for the benefit of another, only makes physiological sense in the context of multicellular populations. In addition to cell death, cannibalism was previously associated with matrix production and, hence, biofilm formation ^20^. However, despite this apparent connection, cannibalism has so far not been studied in the context of differentiated bacterial tissue. We, therefore, initially investigated whether cannibalism mutants of *Bacillus subtilis* derived from the undomesticated, transformable strain 3A38 exhibited altered morphologies in colony architecture compared to the otherwise isogenic wild type strain 3A38 (hereafter referred to as WT). Colony morphology was examined by cultivating strains of interest on agar plates of the biofilm-promoting minimal medium MSgg (minimal salts glycerol glutamate) at 28°C and monitoring biofilm formation over eight days, by capturing daily images (**Fig. 1**). Cannibalistic mutants were either defective in toxin production due to deletions of the structural genes that encode the pre-pro-peptide (*epeX*, *sdpC*, *skfA*), or lacked the autoimmunity functions to the respective toxins (*epeAB*, *sdpI*, *skfEF*). Additionally, we investigated a strain lacking all three cannibalistic toxin-encoding loci (Δ*epeXEPAB* Δ*sdpABCIR* Δ*skfABCEFGH*). This strain is abbreviated as ΔΔΔ and will be referred to as the hypo-cannibalistic mutant.

**Fig. 1.**
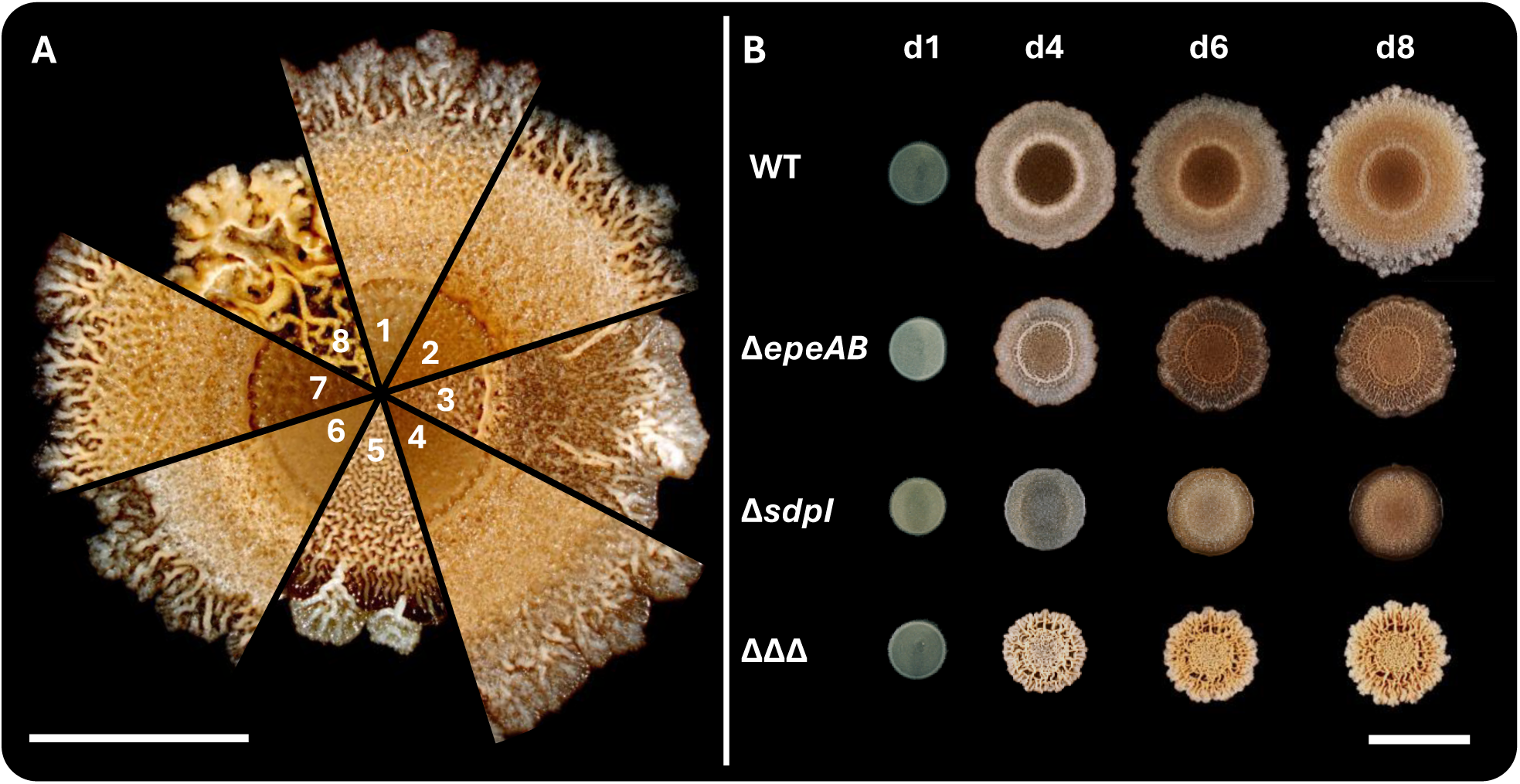
Colony morphologies of *B. subtilis* cannibalistic mutant strains in comparison to the WT. [A] Comparative display of 8 day old biofilms grown on MSgg agar. 1-WT, 2-Δ*epeX,* 3-Δ*epeAB,* 4-Δ*sdpC,* 5-Δ*sdpI,* 6-Δ*skfA,* 7-Δ*skfEF,* 8-ΔΔΔ. Scale bar indicates 5mm. [B] Colony growth and morphology of cannibalistic mutants with distinct phenotypes (Δ*epeAB,* Δ*sdpI*, ΔΔΔ) in comparison to WT over the course of 8 days (day 1 [d1], day 4 [d4], day 6 [d6], day 8 [d8]). Scale bar indicates 10 mm.

The biofilms of all investigated strains displayed a circular symmetry, albeit with different extents of undulating edges. The WT displayed an undulating edge, characterized by vein-like, intertwined, brownish three-dimensional (3D) structures, in agreement with previous descriptions ^3^. The colony exhibited three distinct zones: (i) the center, marking the inoculation site, separated by a ring-like structure from (ii) the inner zone, which was beige, primarily flat, but slightly elevated toward (iii) the outer zone. The outer zone was highly structured, featuring pronounced wrinkles extending to the colony’s edge, with a more brownish-grey hue than the other zones.

Deletion of *epeX*, *sdpC*, or *skfA* had only minor effects on colony morphology, like a somewhat reddish tint across colonies of the *epeX* mutant. The *sdpC* mutant exhibited reduced wrinkle formation in the outer zone, resulting in a flatter, broader surface. Morphological differences in other mutants became apparent from day four onward (**Fig. 1B**). Deletion of the autoimmunity genes for EPE (Δ*epeAB*) or SDP (Δ*sdpI*) led to pronounced changes in colony size, color, and structure. In contrast, Δ*skfEF* exhibited no major effects and retained a WT-like colony appearance (**Fig. 1**, **Fig. S1**). Notably, the *sdpI* (autoimmunity) mutant displayed convoluted structures resembling human brain folds (**Fig. 1A-5**), with an indistinct boundary between the inner and outer zone. Here, the outer zone appeared in a darker brown with reduced structuring. The *epeAB* mutant also showed a color shift toward brown across the entire colony, though all three zones remained distinguishable. Both Δ*epeAB* and Δ*sdpI* biofilms exhibited sporadic white outgrowths in the outer zone, appearing to “escape” from the colony, a phenomenon described in more detail below. The most striking colony architecture occurred in the hypo-cannibalistic mutant (ΔΔΔ). Its biofilm is characterized by a highly wrinkled surface featuring a structurally complex, folded region surrounding the center. The colony displayed increased vertical growth with reduced lateral expansion. Compared to the WT’s beige wrinkles, those of ΔΔΔ were more yellowish, broader, and less defined toward the periphery. In summary, while SKF did not affect the colony structure, the absence of SDP or EPE autoimmunity or – and most pronounced – the complete loss of cannibalistic features (ΔΔΔ) led to significant alterations in colony structure, size, and pigmentation. Cannibalism toxins are, therefore, an important determinant of colony structure, differentiation and expansion.

### Cannibalism toxins show unique and interdependent distribution patterns

Based on the structural consequences of colony architecture described above, we next explored whether specific structural alterations could be correlated with the presence or absence of the three cannibalism toxins within the biofilm. Towards that end, we applied matrix-assisted laser desorption ionization mass spectrometry imaging (MALDI-MSI) ^44^ to visualize the distribution of mature EPE (17 amino acids (aa)), SDP (42 aa), and SKF (26 aa) with a spatial resolution of about 50 µm pixel size. Whole biofilms were grown on 150 µm thick mixed cellulose ester membranes and, following a brief fixation step for inactivation (except for WT strains), were spray-coated with 2,5-dihydroxyacetophenon MALDI matrix (see Materials and Methods for details). A hybrid time-of-flight (QTOF) mass spectrometer (timsTOF fleX MALDI-2, Bruker Daltonics,

Bremen, Germany) was employed to register the MALDI-generated ion traces. MALDI-MSI is a label-free technique that registers all generated ions simultaneously within a mass acquisition range based on their specific mass-to-charge (m/z) values – defined by the molecular composition and charge-providing adduct ion. In our case, the *m/z* values of interest were the following: EPE was recorded most prominently at *m/z* values of 2108.09 and 2146.04 presenting the monoisotopic [M + H]^+^ and [M + K]^+^ ions, respectively; SDP at *m/z* 4314.22 ([M + H]^+^) and 4352.18 ([M + K]^+^), and SKF at *m/z* 2783.31 ([M + H]^+^) and 2821.27 ([M + K]^+^). A sum mass spectrum recorded across a whole WT colony in this *m/z* range is plotted in **Fig. S3A**. All peptides were registered within five ppm of their calculated values. The MALDI-MSI signal intensity was not sufficient to generate MS/MS spectra. A comparison between experimental and calculated ^13^C isotope distributions was used as additional proof of the assignment.. The example of EPE recorded from the WT is depicted in **Fig. S4D**. A series of signals stemming from the acrylate polymer glue used for attaching the biofilm-coated membranes on the glass substrate is moreover notable in the mass spectra. These are found with characteristic mass increments of ethylacrylate units (125.05 Da); acrylate-derived ions of higher abundance are denoted in the mass spectra with asterisks. Zoom-ins into the mass region of the molecular ions, provided in **Fig. S3B** and **Fig. S3C** for the WT (black trace) and selected mutants (red trace, see below for a detailed discussion) show that – if present – all three cannibalism toxins are well detectable with MALDI-MSI method. The recorded ion distribution profiles are again summarized in **Fig. 2A** as a pie chart since all investigated *B. subtilis* biofilms exhibited radial symmetry regarding their morphologies and peptide toxin production. In addition, an MSI overlay of the three toxins showing a whole WT colony is shown in **Fig. 2B**.

**Fig. 2.**
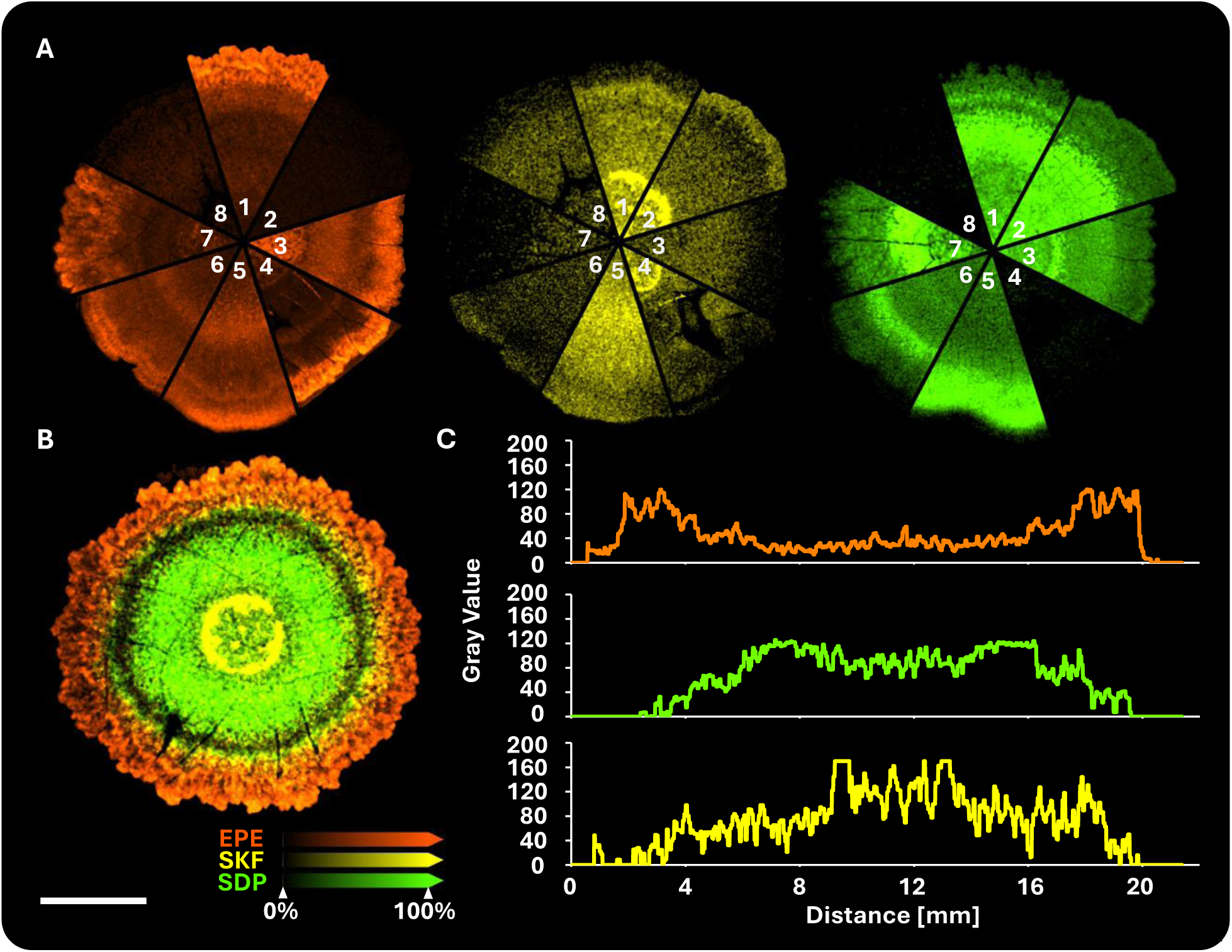
Overlay MALDI-MS images of cannibalistic toxins EPE, SDP and SKF in wild type and mutant *B. subtilis* biofilms. [A] Representative slices of the biofilm covering center, inner and outer zone, displaying the distribution of EPE, SKF, and SDP (from left to right) in the WT (1) and cannibalistic mutant strains (2-Δ*epeX,* 3-Δ*epeAB,* 4-Δ*sdpC,* 5-Δ*sdpI,* 6-Δ*skfA,* 7-Δ*skfEF,* 8-ΔΔΔ). [B] Distribution of EPE, SKF and SDP across the whole colony of the WT. Scale bar indicates 5 mm. [C] 2D-plot of toxin distribution across the distance of the biofilm conducted by linear pixel intensity measurement (gray pixel values). Plateaus at approx. 9-9.7 mm and 12.9-13.2 mm indicate a maximal gray value of 170 in SKF measurement.

EPE (depicted in orange) is predominantly located in the outer zone of the colony (**Fig. 2A-1, 2B, 2C**). As expected, no EPE signal was detected in the *epeX* deletion strain (**Fig. 2A-2**) or the hypo-cannibalistic mutant (**Fig. 2A-8**). The distinct distribution of EPE was lost in the *skfA* mutant, which cannot produce SKF (**Fig. 2A-6**) but slightly accentuated in the *sdpC* mutant, where SDP is absent (**Fig. 2A-4**). The MALDI-MSI data suggest that SKF is particularly enriched in the border region between the central and inner zone of the WT biofilm, forming a distinct ring that correlates well with colony morphology. In the absence of EPE (**Fig. 2A-2**), SKF was “redistributed” to the outer zone, where EPE was “originally” located in the WT. The absence of EPE and SDP-autoimmunity (**Fig. 2A-3/5**, respectively) resulted in the loss of the characteristic SKF ring in the colony center, leaving a weak and broad distribution pattern across the remaining biofilm. No SKF was detected in the *skfA* deletion strain or the hypo-cannibalistic mutant, confirming the absence of EPE, SDP, and SKF in this mutant. Furthermore, SKF was undetectable in the *skfEF* mutant strain (**Fig. 2A-7**, **Fig. S3**), suggesting that SKF production depends on the presence of its respective autoimmunity factors. Such a connection between autoimmunity and production has not yet been observed, but is a unique feature of SKF that stands in contrast to SDP and EPE. While the false-color images indicate a weak signal throughout the biofilm, the corresponding mass spectra revealed that these base signals represent noise rather than the presence of SKF (**Fig. 2A-6/7/8**, **Fig. S3**).

**Fig. 3.**
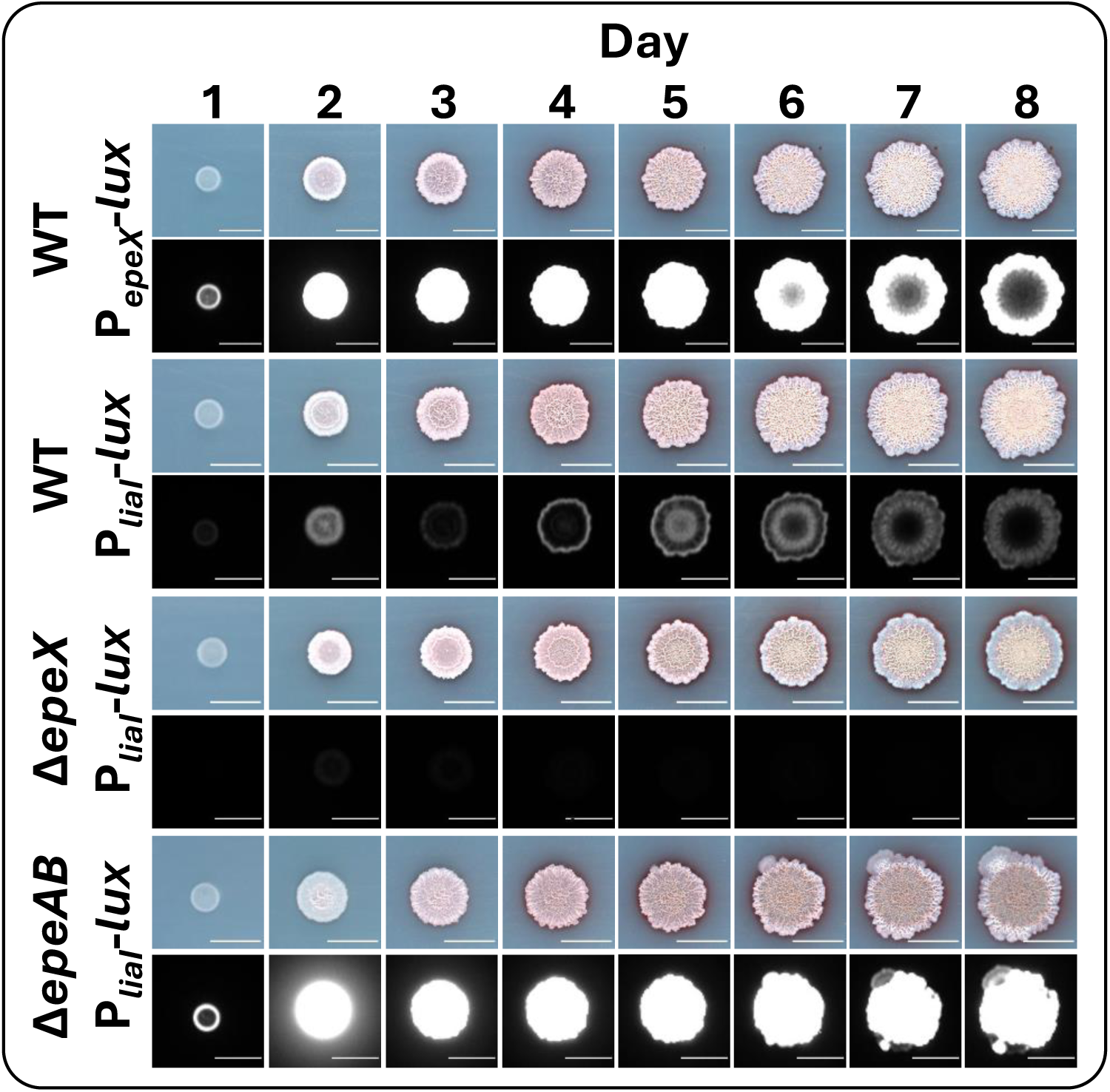
Activity of promoters of *epeX* and the corresponding stress response *liaI*. fused to a *luxABCDE*-cassette in WT and mutant strains on MSgg agar. Colony expansion and luminescence signal distribution are shown over the course of 8 days. Scale bar indicates 10 mm.

**Fig. 4.**
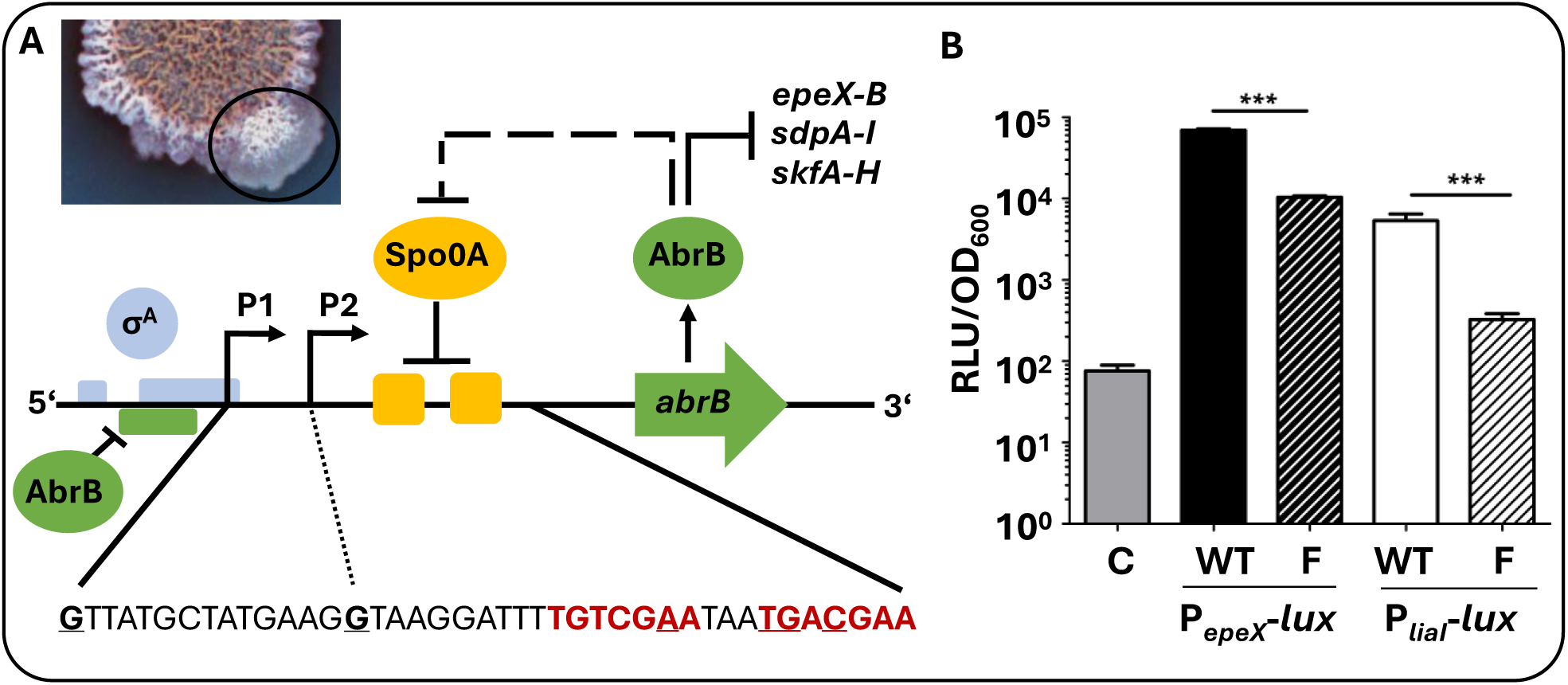
Various, singularly occurring single nucleotide polymorphisms (SNPs) in the Spo0A binding sites of the *abrB* promoter region allows escape mechanism in suppressor mutants of Δ*epeAB* strains. [A] Image of an Δ*epeAB* colony with a circle highlighting the outgrowth of a flare (suppressor mutant). Beneath, a scheme showing the location of the SNPs within the Spo0A binding sites (red underlined nucleotides) and the influence they have on the intricate interplay of Spo0A and AbrB in regard to cannibalistic operon expression. [B] Significant differences in activity of P*_epeX_*-lux (black) and P*_liaI_*-lux (white) in the wild type (WT) and Δ*epeAB* flares (F) showing a reduced expression of *epeX* and the corresponding stress response *liaI* in those suppressor mutants (*** indicate a P value <0.001). C indicates the control (P*_empty_-lux*).

In the WT biofilm, SDP (depicted in green) was primarily located in the inner zone (**Fig. 2A-1**). However, its distribution is significantly altered in strains lacking the autoimmunity genes for EPE (Δ*epeAB*, **Fig. 2A-3**) or SDP (Δ*sdpI*, **Fig. 2A-5**), as well as in the absence of SKF. Within the biofilm of the *epeAB* mutant, SDP strongly accumulates at the central ring structure, whereas in a *sdpI* mutant biofilm, SDP is predominantly found in the outer zone. In the *skfA* mutant, SDP localizes strongly at the border between the inner and outer zones of the biofilm. Interestingly, the distribution of SDP in a *skfEF* mutant is comparable to the WT, even though we demonstrated that no SKF is in this mutant. Hence, this effect might depend on the SkfEF function rather than SKF itself. As anticipated, SDP was not detected in the *sdpC* deletion and the hypo-cannibalistic mutant.

**Fig. 5.**
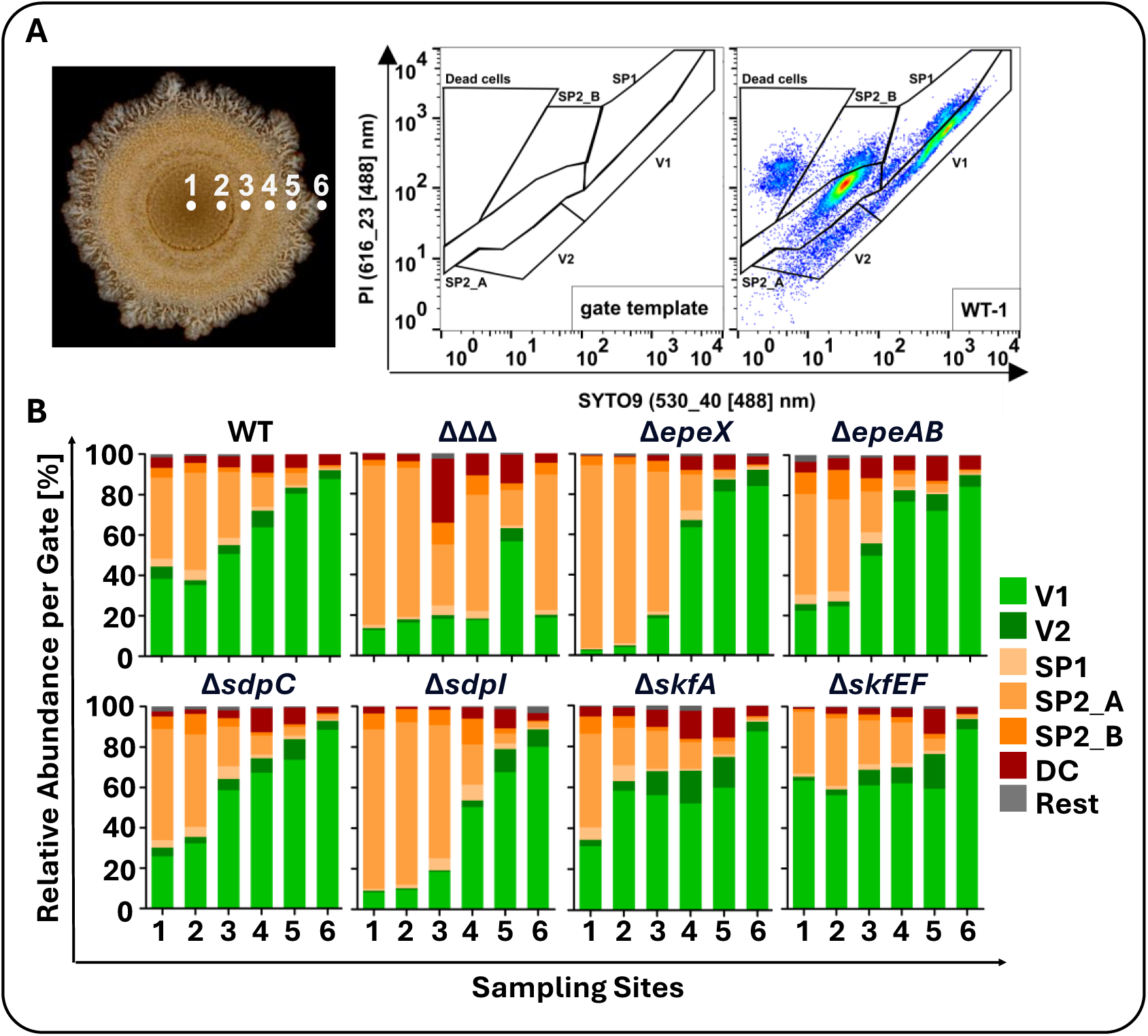
Distribution of live, dead and sporulating cells across *B. subtilis* colonies in regard to cannibalism. [A] Sampling sites across an exemplary WT colony are shown, as well as the gate template for analysis and sorting together with an exemplary dataset of flow cytometric measurement of cells present in sampling site 1 of the WT. PI signal is plotted against SYTO9 signal. [B] Overview of flow cytometric analysis results are summarized in bar graphs for each strain, where the relative abundance of cells per gate are plotted against sampling sites. V1 and V2 indicate vegetative cells, SP1 – SP2_B represent different spore types, DC indicates dead cells.

The combined picture of the distribution of the three toxins in the WT strain reveals that they collectively cover the entire colony area, each with its preferred zone (**Fig. 2B**): A quantitative analysis of the gray pixel values in the WT biofilm was conducted by linear pixel intensity measurement (**Fig. 2C**). Concurrent with the afore-described MALDI-MSI results, EPE predominantly covers the outer zone, SDP is the inner zone, and SKF peaks in the center of the colony, although it is also lightly present in other zones. This suggests a distributional interdependence among the three toxins that requires further investigation.

Remarkably, a closer look into the peptide mass range revealed that, especially for hypo-cannibalistic ΔΔΔ, an entire set of newly expressed biomolecules is detectable (**Fig. S4**), which strongly correlates with the colony structure. The example of the ion registered at *m/z* 2143.23 is shown in **Fig. S4C**. Future work will address these findings and the possible identity of these novel compounds in more detail. These preliminary results nicely underscore the analytical potential of the MALDI-MSI technology as a discovery tool.

### Expression of *epeX* correlates with EPE production

We had previously demonstrated that cannibalism toxins intrinsically trigger the CESR ^24,31^. P*_liaI_*, P*_bceA_*, and P*_psdA_*, when fused to the *luxABCDE* reporter cassette, provide a highly specific, cannibalism-inducible luminescent readout, with P*_liaI_* responding to EPE stress. At the same time, P*_bceA_*/P*_psdA_* were triggered by both SDP and, particularly, by SKF, at least in liquid cultures of strains derived from the domesticated laboratory reference strain W168. We, therefore, tested if these reporters could also be applied to record cannibalism stress during colony development in undomesticated strains. The spatiotemporal distribution of toxin gene expression (P*_epeX_*, P*_sdpA_*, and P*_skfA_* for EPE, SDP, and SKF, respectively) and the corresponding CESR in developing biofilms was monitored over the course of eight days. Promoter activities were examined in WT biofilms and autoimmunity-deficient strains for all three toxins. For SKF and SDP, the results obtained for W168-derived reporters in liquid culture could, unfortunately, not be reproduced for colonies of 3A38-derived strains. We therefore focused on the two promoters associated with EPE-related stress (**Fig. 3**).

The promoter driving *epeX* expression, P*_epeX_*, exhibited vigorous activity throughout the whole developmental period, with a prominent signal observed in the outer zone of the biofilm, resembling the inoculation spot, already on day 1 (24 hours post-inoculation). From day 2 to day 5, the signal spread across the entire colony. From day 6 onwards, activity steadily decreased from the colony’s center so that by day 8, P*_epeX_* activity was confined to the outer zone. The corresponding EPE stress-responsive promoter, P*_liaI_*, revealed an intriguing ring pattern during biofilm growth. However, from the third day onwards, the P*_liaI_* signal was predominantly located at the edge of the biofilm and – particularly towards the end of colony differentiation, the activity of P*_epeX_* and P*_liaI_* co-localize (days 7-8). The distribution of the P*_liaI_-lux* signal indicates that, while the *epeXEPAB* locus is consistently expressed (P*_epeX_-lux* readout), active EPE is not directly released.

This observation indicates that the maturation or release of active EPE is tightly regulated beyond the transcriptional control of P*_epeX_*. Importantly, the P*_liaI_-lux* readout at the end of colony development perfectly overlaps with the location of mature EPE toxin, as detected by mass spec imaging (**Fig. 2**). Since P*_liaI_-lux* seemed indeed suitable as a readout for EPE release, we next probed P*_liaI_* activity in an EPE-OFF strain (Δ*epeX*). The absence of any P*_liaI_* activity in this strain corroborates that the Lia system specifically responds to EPE stress in differentiating colonies. In contrast, the massively increased P*_liaI_-lux* signal in the autoimmunity mutant (Δ*epeAB*) demonstrates that EpeAB indeed counteracts EPE stress in colony biofilms. Without this ABC transporter, P*_liaI_* activity was persistently strong and spread across the entire colony, resembling the pattern seen in the WT for P*_epeX_*.

### EPE stress in autoimmunity mutants gives rise to suppressor mutants

Interestingly, from day 7 onward, only weak P*_liaI_* activity was detected in specific areas of the colony edge, indicating a reduction in EPE stress due to decreased EPE production. Moreover, these “flares” remained white, suggesting a lack of sporulation. Such outgrowths were frequently observed at the edges of biofilms from strains harboring either *epeAB* or *sdpI* deletions. When isolated, their white color and altered colony morphology remained segregationally stable (**Fig. S2**), indicating that these flares represent suppressor mutants that occurred under severe cannibalism stress. One of these isolated spontaneous mutants was subsequently characterized by whole-genome sequencing of genomic DNA and luminescence-based analysis of P*_epeX_* and P*_liaI_* promoter activities in the corresponding strains. We identified a single nucleotide polymorphism (SNP) in the Spo0A binding site within the promoter region of *abrB* (**Fig. 4A**).

Sequence analysis of the *abrB* promoter region from the other flares identified additional – but not identical – SNPs in the Spo0A consensus sequence (TGNCGAA) at genome positions 3,508,948 (G to A), 3,508,949 (T to C), 3,508,954 (A to G), and 3,508,946 (C to T).

It is known that Spo0A binding to this promoter results in a repression of *abrB* expression ^16^. Interferences with Spo0A binding should result in elevated AbrB levels and, hence, a repression of its target genes, thereby suppressing sporulation but also cannibalism toxin production. This is precisely what we observed when studying the respective target promoters. The G to A substitution in the Spo0A binding site of the *abrB* promoter identified in flare suppressor mutant 1 affected not only biofilm morphology and sporulation (**Fig. 4**) but also *epeX* and *sdpC* expression negatively, as a consequence of *abrB* derepression (**Fig. S2**). As a consequence of the approximately 10-fold reduction in P*_epeX_* activity, the corresponding EPE stress response in the flare mutants, monitored by P*_liaI_* activity, was also reduced by a factor of 10 compared to the WT. This result indicates that the EPE- or SDP stress is so severe within the biofilm context that it gives rise to suppressor mutants that interfere with the central stationary phase regulation. The transition state regulator AbrB controls over 250 genes during the transition between vegetative growth and the onset of stationary phase, including functions involved in sporulation, motility, matrix formation, quorum sensing, and cannibalism toxin production ^45–47^. The derepression of *abrB* is, therefore, a powerful mechanism for escaping cannibalism stress, as induced by EPE, through the suppression of toxin production based on inhibition of Spo0A binding at the *abrB* promoter.

### Cannibalism toxins affect cell type distribution within biofilms

The architecture of multicellular colony biofilms is the consequence of a spatiotemporal controlled phenotypic diversification of different cell types – a result of the regulatory cascades governing differentiation in response to external stimuli such as nutrient and oxygen availability in combination with the underlying stochastic gene expression ‘noise’ ^48–53^. Such spatial patterning has also been observed for *B. subtilis* colonies: Matrix-producing cells are located primarily in the upper layers of the biofilm, providing structural integrity through the secretion of extracellular components. Motile cells are found at the periphery, where swarming motility facilitates expansion. Sporulating cells are found in nutrient-deprived central-upper regions of the colony, which partially coincides with cannibalistic cells that delay spore formation by lysing sibling cells for further nutrient supply. Finally, competent cells, which can take up extracellular DNA to enhance genetic diversity, are dispersed heterogeneously ^3,6,54^. Given the observed influence of cannibalism toxins on colony architecture and their interdependent distribution patterns, we wondered whether the cell type distribution within colonies was also affected as a function of cannibalism toxin action. We, therefore, investigated the spatial occurrence of different cell types throughout the colony. Towards this goal, we employed colony biopsy in combination with microbial flow cytometry and cytometric fingerprinting ^43^ on fully grown biofilms (8 days old). Biopsies were collected from six different sampling sites of the biofilm, covering zones with structural differences as well as areas with distinct toxin distributions (**Fig. 5A**). Each sample was then analyzed for the presence of spores, live and dead cells (**Fig. 5**) as well as cell cycle states, to assess population heterogeneity (**Fig. S6**). Both intrinsic parameters, such as scattering, and extrinsic parameters, such as fluorescence, were measured using flow cytometry. To differentiate living from dead cells, a combination of SYTO9 (a green-fluorescent, membrane-permeant nucleic acid stain) and propidium iodide (PI, which indicates compromised cell membranes and cell walls) was used for staining (**Fig. 5**). In addition to the SYTO9-/PI- staining, cell cycle states were assessed using 4’,6-di-amidino-2-phenylindole (DAPI), a dye that intercalates into AT-rich regions of double-stranded DNA. The resulting cell subsets are displayed as 2D plots, with cell gates defined for fluorescence-activated cell sorting (FACS) (**Fig. S5**, **Fig. S7**). Based on these gates, we analyzed the changes in cell type distribution between the WT and cannibalism mutant strains.

Six distinct subpopulations could be identified within *B. subtilis* biofilms, based on SYTO9-/PI- staining: three spore types (SP1, SP2_A, SP2_B), two vegetative cell types (V1, V2) and dead cells (DC). Dead cells exhibited only red PI fluorescence, while spores showed low to medium SYTO9 fluorescence compared to vegetative cells, which appeared in the rightmost gates of the 2D plots (**Fig. 5A**). In the WT, a high percentage of spores were detected at sampling sites 1-3 (48.9%, 58.1%, 38.5%, respectively) (**Fig. 5B**, **Fig. S6**), located in the center and the adjacent area of the inner zone. Within the inner zone (sampling site 4), the percentage of spores decreased to 18.7%, while vegetative cells increased to 72%. This trend continued towards the outer zone (sampling sites 5 and 6), where only 2.5% of cells were spores, but 92% were vegetative cells, with a moderate 5% of dead cells. The analysis of DAPI-stained cells (**Fig. S5**, **Fig. S7**) supported this distribution, with an excess of dormant spores and sporulating cells found in the center of the colony (53.69% at sampling site 1), and the proportion of spores also decreasing towards the outer zone of the biofilm (sampling site 6).

This cell type distribution was affected in most cannibalistic mutants. Specifically, the Δ*epeX*, Δ*epeAB*, and Δ*sdpI* strains showed a noteworthy increase in spore levels at sites 1-3, while Δ*sdpC*, Δ*skfA*, and Δ*skfEF* mutants displayed results like the WT. The increased spore abundance, particularly in the center and adjacent regions of the inner zone, may explain the red coloration observed in the *epeX* mutant. Previous studies have shown that sites of sporulation tend to exhibit darker biofilm coloration ^55,56^. The rise in spore numbers in toxin deletion strains is consistent with earlier observations ^20^, further supporting the hypothesis that cannibalism is a strategy to delay sporulation. Interestingly, this pattern was only observed in the *epeX* and *sdpC* mutants but not in the *skfA* mutant for both the SYTO9-PI and the DAPI-staining analyses (**Fig. 5**, **Fig. S5**), again emphasizing that SKF does not seem to play a major role in the cannibalism-dependent structuring of colony biofilms.

Based on the increased CESR in autoimmunity-deficient mutants, we expected an increase in dead cells in these strains. However, the flow cytometry data did not support this assumption. While approximately 5.6% of cells were dead in the WT colony, only a slight increase was observed in the *epeAB* mutant (8.4%), while the abundance of dead cells across the entire colony decreased in the *sdpI* (3.8%) and *skfEF* (4.3%) mutants.

Again, the most remarkable results came from the hypo-cannibalistic mutant, which was also most severely affected in its colony architecture (**Fig. 1**). Colony biopsy and flow cytometry revealed a completely rearranged distribution of vegetative, dead, and sporulating cells in comparison to the WT. In this hypo-cannibalistic mutant, spores were highly abundant across all sampling sites, even in the outer zone. The entire colony of the cannibalistic triple mutant contained 58% spores and only 34% vegetative cells, whereas the WT had 22% spores and 71% vegetative cells. Comparable results were obtained in the DAPI analyses (**Fig. S5**, **Fig. S7**). While the overall abundance of dead cells increased only slightly by 0.94% in the triple mutant compared to the WT, a closer examination of the spatial distribution of dead cells revealed significantly elevated percentages at sampling sites 3-5 (31.6%, 10.6%, 14.2%, respectively). This pattern was not observed in any of the other mutants and may offer an explanation for the highly folded structure of the colony.

Taken together, the flow cytometry results are in good agreement with the effects on colony architecture observed in the different mutant strains. The more severely the overall structure was affected, the more pronounced the underlying changes in cell type distribution throughout the colony. This is particularly pronounced in the hypo-cannibalistic mutant, which shows a complete loss of the zoning in its colony compared to the WT: The colony does not expand much beyond the area initially spotted but instead wrinkles uniformly across its complete surface (**Fig. 1**). This is reflected by the even distribution of spores through the colony (**Fig. 5B**). The combined action of the three cannibalism toxins is crucial for both structure and function of the differentiated colonies, presumably by affecting the spatiotemporal control of the underlying processes.

## Discussion

*B. subtilis* has evolved multiple survival strategies in response to the perpetual shifts in environmental conditions, and to compete with other organisms for nutrients. Differentiation in the context of multicellular aggregates such as colony biofilms gives rise to the development of different specialized cell types. This division of labor enhances the adaptability of an organism to its environment and thereby increases the chances for survival, particularly in unpredictable habitats, where change is the only constant ^57^. In *B. subtilis,* a complex regulatory network orchestrates and coordinates these differentiation strategies and ensures an appropriate adaptation. Under conditions of prolonged nutrient deprivation, cells of *B. subtilis* can initiate sporulation, which will ultimately result in dormant and highly resistant endospores that can outlast long periods of adverse conditions while maintaining the ability to germinate once the conditions become more favorable again ^58^. Since sporulation is an irreversible process and both time- and energy-consuming, *B. subtilis* employs other survival strategies aimed at preventing or at least delaying commitment to this strategy of last resort.

Cannibalism has been described as a survival mechanism that delays sporulation under nutrient-limiting conditions by enabling a subpopulation of cells to lyse their neighbors and utilize the released nutrients ^23^. Earlier studies already demonstrated the cannibalism-dependent delay of sporulation for the domesticated laboratory strain: in *sdp/skf* mutants, heat-resistant colony-forming units appeared earlier and in higher numbers compared to the wild type ^20,21^. In the present study, we verified this role of cannibalism in the context of differentiated colony biofilms of strain 3A38, a transformable derivative of the undomesticated ancestor strain: the hypo-cannibalistic mutant (ΔΔΔ, **Fig. 5**), which is unable to produce any of the three cannibalistic toxins, showed a substantial increase in spore abundance across the whole colony. In contrast to the initial studies cited above, which only considered SDP and SKF, we conclude that EPE and SDP, but not SKF, are primarily important in delaying the onset of sporulation since the *skfA* mutant does not display a higher spore abundance compared to the WT (**Fig. 5**).

While delaying sporulation is, therefore, indeed, an important function of cannibalism toxins, irrespective of different strain backgrounds and physiological settings, the mechanism by which this is achieved seems to differ between them. The initial hypothesis suggested that the altruistic killing of one subpopulation by another is the reason for the sporulation delay due to the released nutrients ^23^. In contrast, our current study did not provide any evidence for cannibalism-dependent killing, at least not in colony biofilms. Instead, cell death seems primarily associated with sporulation itself since the number of dead cells increases with the sporulation rate but is independent of the presence or absence of cannibalism toxins (**Fig. 5**).

Instead of cannibalism-induced lysis, we could demonstrate a crucial role of cannibalism toxins for structuring and functionalizing colony biofilms (**Fig. 1**). While some indirect evidence for a link between SDP/SKF and matrix production had indicated a role of cannibalism toxin in biofilm formation ^20^, a direct impact of cannibalism on multicellular differentiation had so far not been reported. Cannibalism affects both the 3D-structuring of colonies but also balances differentiation and colony expansion, as most prominently observed for the hypo-cannibalism strain (**Fig. 1**). Colonies of this cannibalism-deficient strain were characterized by hyper-wrinkling, a lack of colony expansion and a massively increased sporulation rate throughout the colony. This result suggests that cannibalism toxins balance colony differentiation by restricting the degree and amount of wrinkling (growth in height), thereby ensuring that resources are provided for colony expansion (growth in width). This hypothesis would also explain the results from colony biopsy and flow cytometry. Growing in height moves a significant fraction of cells in the multicellular tissue away from the agar surface and, hence, away from the source of nutrients. Sporulation and cell death would be the consequences, as observed for the hypo-cannibalism strain (**Fig. 5**). In contrast, colony expansion increases the colony surface area that come into contact with the substratum, thereby providing access to new sources of nutrients.

In the context of colony differentiation, the absence of autoimmunity against EPE/SDP – but not SKF! – had the strongest impact of individual mutations on colony morphology (**Fig. 1**) and the distribution and abundance of the distribution of different cell types across the colony biofilm (**Fig. 5**). Moreover, we observed a massively increased CESR in colonies lacking the autoimmunity at least for EPE (**Fig. 3**). The resulting stress seemed to provoke the generation of suppressor mutations to circumvent the damage inflicted upon the cells (**Fig. 4**), which highlights the importance of a functional defense system against intrinsically produced toxins. While we did not detect any increased cannibalism-dependent cell death under these conditions, we did observe a strongly increased LiaRS-dependent stress in colonies lacking the EPE autoimmunity (**Fig. 3**). Our results also shed new light on the sporulation killing factor, SKF. The eponymous killing property of SKF was so far only demonstrated in liquid cultures ^23^. Under these conditions, we also observed a strong CESR based on the P*_psdA_* and P*_bceA_* biosensor read-out ^24^. In contrast, we could not observe any inhibitory activity of SKF for colony biofilms grown on agar plates, and no CESR could be detected with the same reporter strains as above. This could be attributed to either the inability to adequately enrich SKF within the plate-based assays, or the explanation may be found in the distribution of the toxin in the surface-attached biofilm. Our MALDI-MS imaging experiments demonstrated that SKF is predominantly located in the center of the biofilm and, to a lesser extent, at the periphery. The latter finding is consistent with an independent study demonstrating the localization of SKF at the outer boundary of the biofilm and additionally showing, also employing MALDI-MS imaging, that a *srfAC* mutant, which could no longer produce surfactin, was deficient in SKF production ^59^. Cannibalism has previously already been associated with surfactin ^20^. Considering these recent findings, this provokes the question of how and why this biosurfactant co-occurs with the cannibalism toxin SKF at the outer zone of the biofilm. For surfactin, the localization can be explained due to its surface tension-lowering properties, which also facilitate the spreading and expansion of multicellular colonies on the substrate ^60^. For SKF, this co-localization raises the question of whether this peptide might also act as a surface-active compound involved in biofilm expansion rather than killing for nutrients. This new perspective on the role of SKF is further supported by preliminary data on the three-dimensional resolution of toxin distribution in thin sections of *B. subtilis* biofilms, where SKF is predominantly found underneath the biofilm, that is, at the interphase to the agar surface (personal communication K. Dreisewerd, ^61^) (**Fig. 6**). This finding demonstrates the considerable potential of resolving toxin distribution in three dimensions. Moreover, it highlights the need for continued advancement in the field, notwithstanding the already substantial progress in methodology and technology in recent decades ^41,44,62,63^. Further development of the methodology for investigating bacterial biofilms will significantly enhance our understanding of the localization of the relevant biomolecules.

**Fig. 6.**
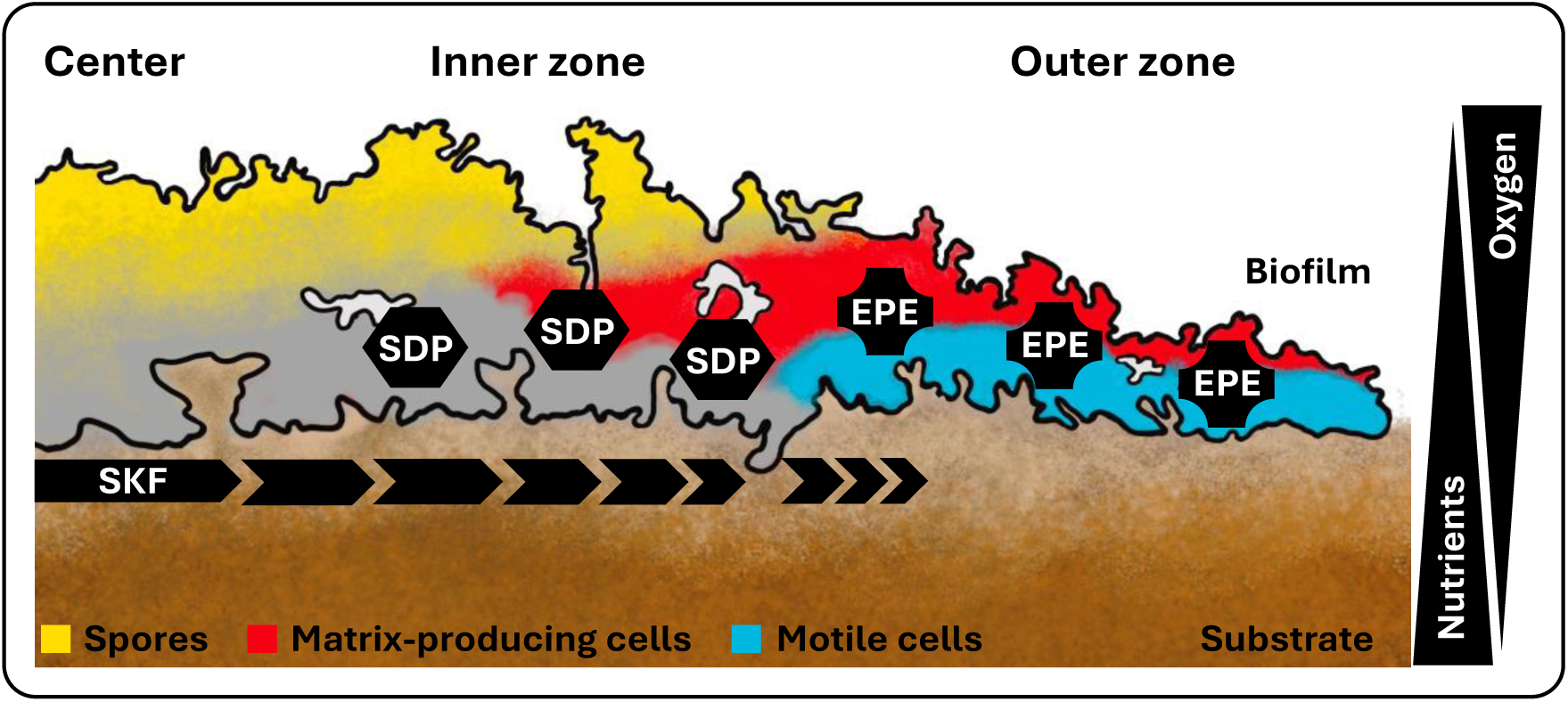
Schematic illustration of a *B. subtilis* biofilm cross section showing the distribution of differentiated cell types and cannibalistic toxins SKF, SDP, and EPE across the biofilm. Based on Vlamakis et al., 2008 ^54^.

Nevertheless, state-of-the-art MALDI-MS imaging technology has enabled us to analyze the distribution of EPE, SDP, and SKF across biofilms of *B. subtilis* and cannibalistic mutants at high spatial resolution. This allowed gaining a comprehensive picture of the distribution of cannibalism toxins (in 2D) and their changes as a function of genetic manipulations of cannibalism genes. As mentioned before, SKF is concentrated in the center of the biofilm, with some minor occurrence at the outer edges of the colonies, while SDP is present in the inner zone of the biofilm (**Fig. 2**). A similar pattern of SKF and SDP distribution has previously been reported for *B. subtilis* biofilms grown on MSgg agar by using MALDI-MS imaging ^64^. In a third study, the differences in SDP/SKF localization can be attributed to the difference in strains and culture conditions applied: the authors investigated the domesticated strain *B. subtilis* PY79 grown on ISP2 medium, which lead to relatively unstructured biofilms ^65^. The present study builds upon and complements the knowledge generated by these previous studies by adding the EPE toxin to the picture and demonstrating the co-occurrence and interdependence with the other two toxins. The epipeptide EPE is found predominantly within the biofilm’s outer, actively growing zone (**Fig. 2**). It is known to trigger the CESR and to cause cell death via massive membrane perturbations ^22,29,66^.

Our study demonstrates the importance of cannibalism for the structural organization of *Bacillus* biofilms. As we have shown, toxin production is tightly linked to wrinkle formation and colony expansion. While the absence of one toxin can be compensated for by the presence of the other two toxins, so that individual mutants in *epeX, sdpC*, nor *skfA* did not display strong phenotypes, the absence of all three toxins at the same time dramatically influences the structure and shape of the biofilm (**Fig. 1**). Based on our findings and the aspects discussed above, we propose a modified model for cannibalism in *B. subtilis* multicellular life (**Fig. 6**).

Cannibalism acts as a differentiation strategy that delays the commitment to sporulation. The absence of cannibalism toxins results in hyper-sporulating, non-expanding colonies dominated by spores with few vegetative cells. Endospore formation involves mother cell lysis, increasing dead cell abundance in a hypo-cannibalistic environment. The accumulation of dead cells induces mechanical buckling, promoting vertical biofilm growth ^67^, which in turn drives sporulation further as cells are pushed away from nutrient sources. In the presence of EPE, SDP, and SKF, hyper-sporulation is prevented by a fine-tuned system of localized PCD, allowing susceptible sibling cells to be sacrificed progressively and simultaneously across the biofilm. This results in structuring the colony so that the nutrients that are provided allow the population to thrive further, thereby also reaching new sources of nutrients at the periphery of the expanding colony. This process ensures lateral colony expansion while suppressing vertical buckling, thereby, also restricting sporulation. Together, this allows *B. subtilis* colonies to grow and thrive on agar surfaces.

Production of the ECM is linked to nutrient availability, mainly on the propagating front of the biofilm but also close to the substrate in older parts of the colony ^68^. These findings are corroborated by our data concerning the localization of cannibalism toxins: EPE is found at the periphery of the developing biofilm, potentially controlling colony expansion. SKF seems to be located at the interface between the biofilm and the substrate (personal communication, K. Dreisewerd, ^61^) and may help colony expansion in conjunction with surfactin.

Our findings suggest that cannibalism is not merely a survival strategy to delay sporulation, but also a key mechanism to control biofilm development and colony expansion in *B. subtilis*. The interplay between toxin production and differentiation ensures a dynamic and organized colony structure. Future research must address the spatiotemporal correlation between swarming, matrix production, and sporulation as a function of cannibalism toxin production. While our study sheds light on the structural role of cannibalism and its functional consequences on the phenotypic and cell type distribution within colony biofilms, it also stresses the importance of bacterial PCD for multicellular differentiation and the structuring of bacterial tissue – a connection that is already well-established for higher eukaryotic tissue but has only emerged recently for bacteria, owing to recent technological breakthroughs that finally allow resolving bacterial macroscopic structures at or at least near single-cell resolution ^36^. Studying bacterial tissue formation has just begun, but is finally achievable.

## Materials and Methods

### Reagents

Chemicals used in this study were obtained from *Thermo Scientific* (Waltham, MA, USA), *Carl Roth GmbH & Co. KG* (Karlsruhe, Germany) or *Sigma-Aldrich* (St. Louis, MO, USA), if not otherwise stated. All enzymes (restriction endonucleases, ligases and polymerases for PCR) originated from *New England Biolabs* (Ipswich, MA, USA) and general cloning procedures were performed according to the recommended protocols. PCR purifications and plasmid preparations were carried out using the corresponding kits from *Süd-Laborbedarf GmbH* (Gauting, Germany).

### DNA isolation

#### Isolation of genomic DNA from B. subtilis for transformation

800 µL of overnight culture was transferred into a 2 mL reaction tube and mixed with an equal volume of SC buffer (NaCl 0.15 M, sodium citrate 0.01 M, pH 7.0). Cells were then harvested by centrifugation at 13,300 rpm for 1 min and afterwards resuspended in 1 mL SC buffer. 20 µL lysozyme (15 mg/ml) was added and the mixture was incubated at 37 °C for 15 min. Following cell lysis, 800 µL of 5 M NaCl and 200 µL H_2_O were added, and the mixture was subsequently filtered through a 0,45 µm syringe filter (*Sarstedt AG & Co. KG*, Nümbrecht, Germany). The DNA solution was directly used for the transformation of *B. subtilis* or stored for later use at −20 °C.

#### Isolation of total DNA from *B. subtilis*

For isolation and rapid purification of highly pure genomic DNA of *B. subtilis*, the NucleoSpin Microbial DNA (*Macherey-Nagel GmbH & Co. KG,* Düren, Germany) kit was used according to the manufacturer’s instructions.

### Strain construction

All oligonucleotides and strains used in this study are listed in **Table 1** and **Table 2**, respectively. The synthesis of oligonucleotides was performed by *Microsynth AG* (Balgach, Switzerland). DNA sequencing was performed by *Eurofins Genomics Europe Shared Services GmbH* (Ebersberg, Germany).

**Table 1.**
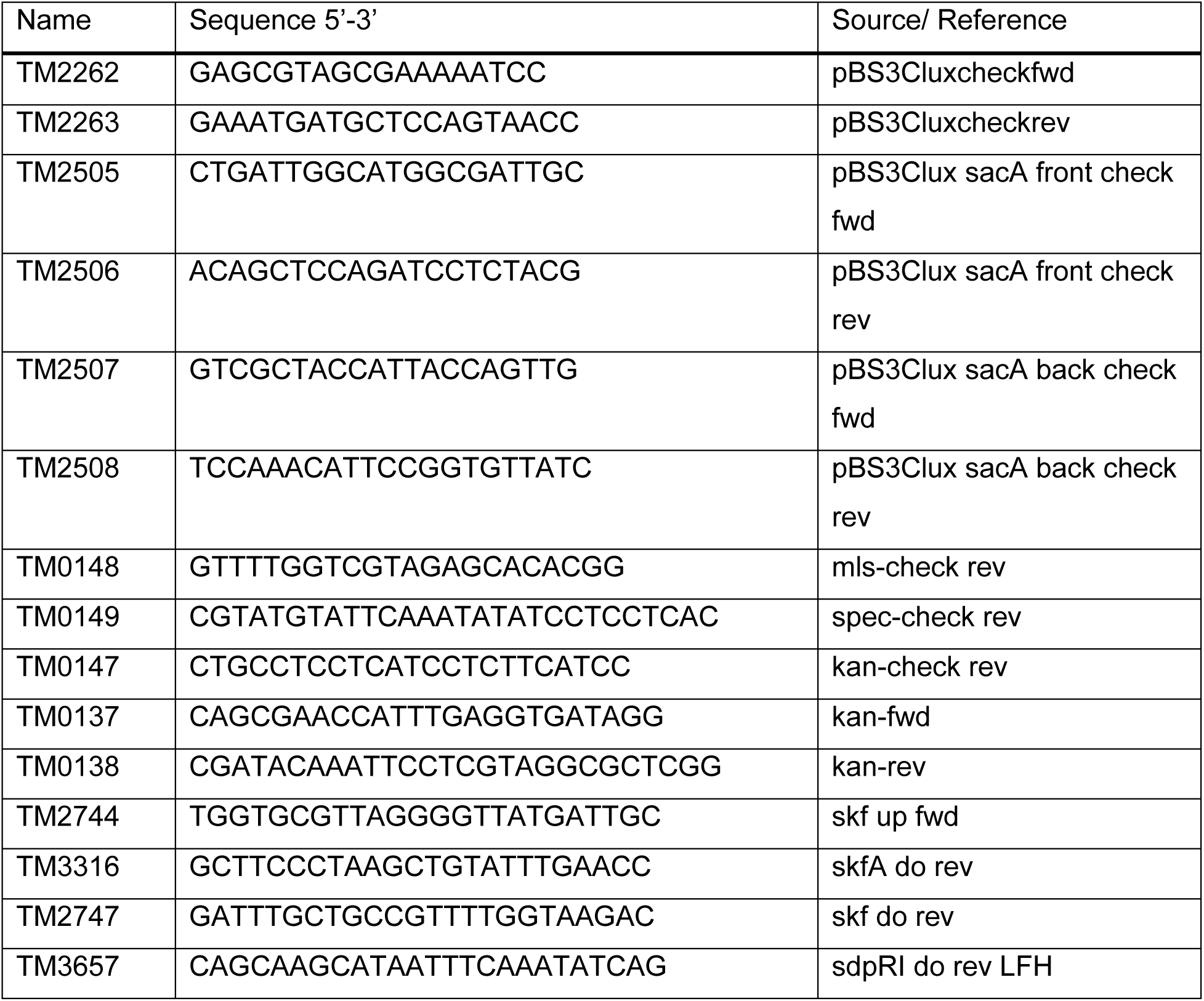
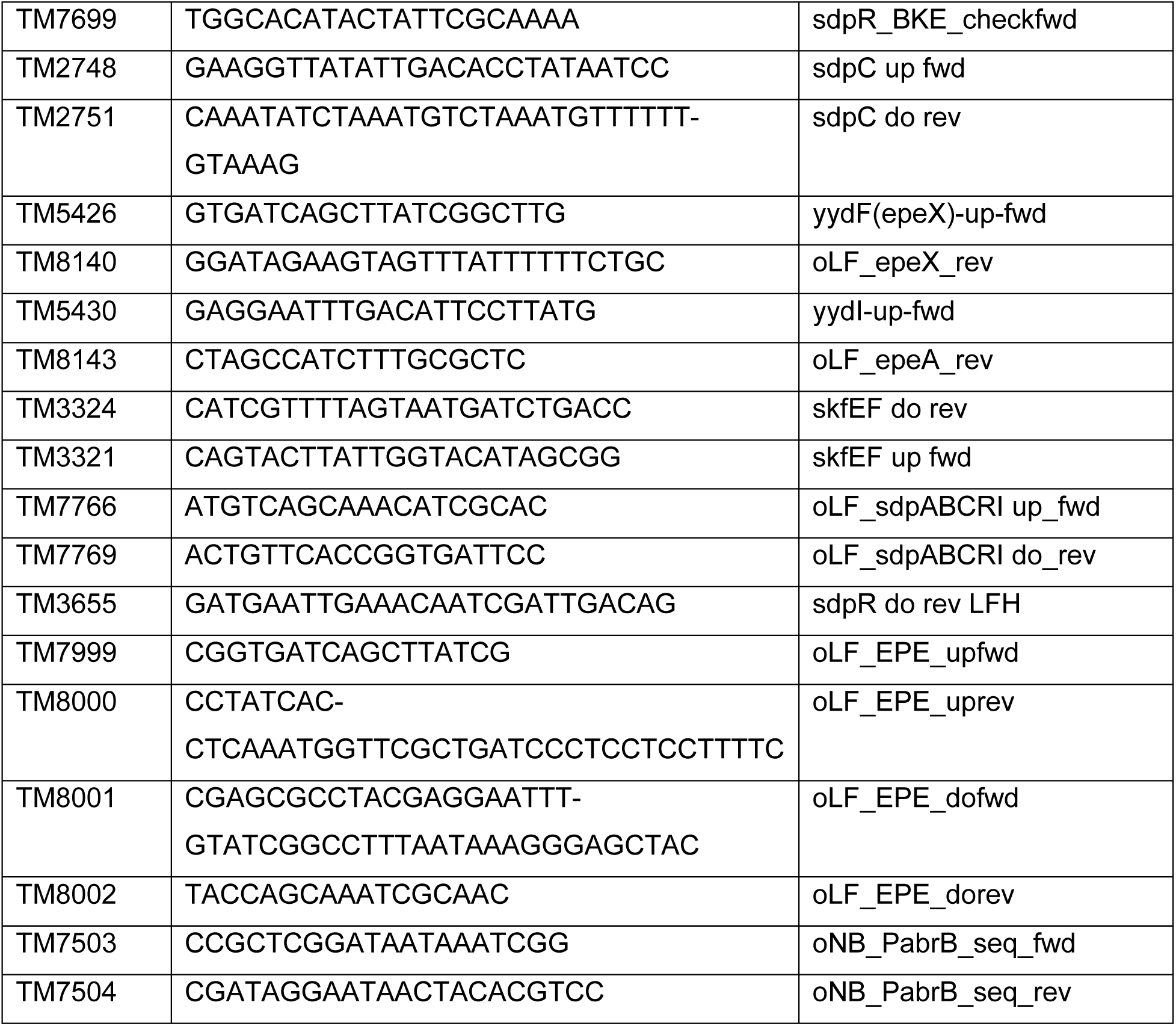
Oligonucleotides used in this study.

**Table 2.**
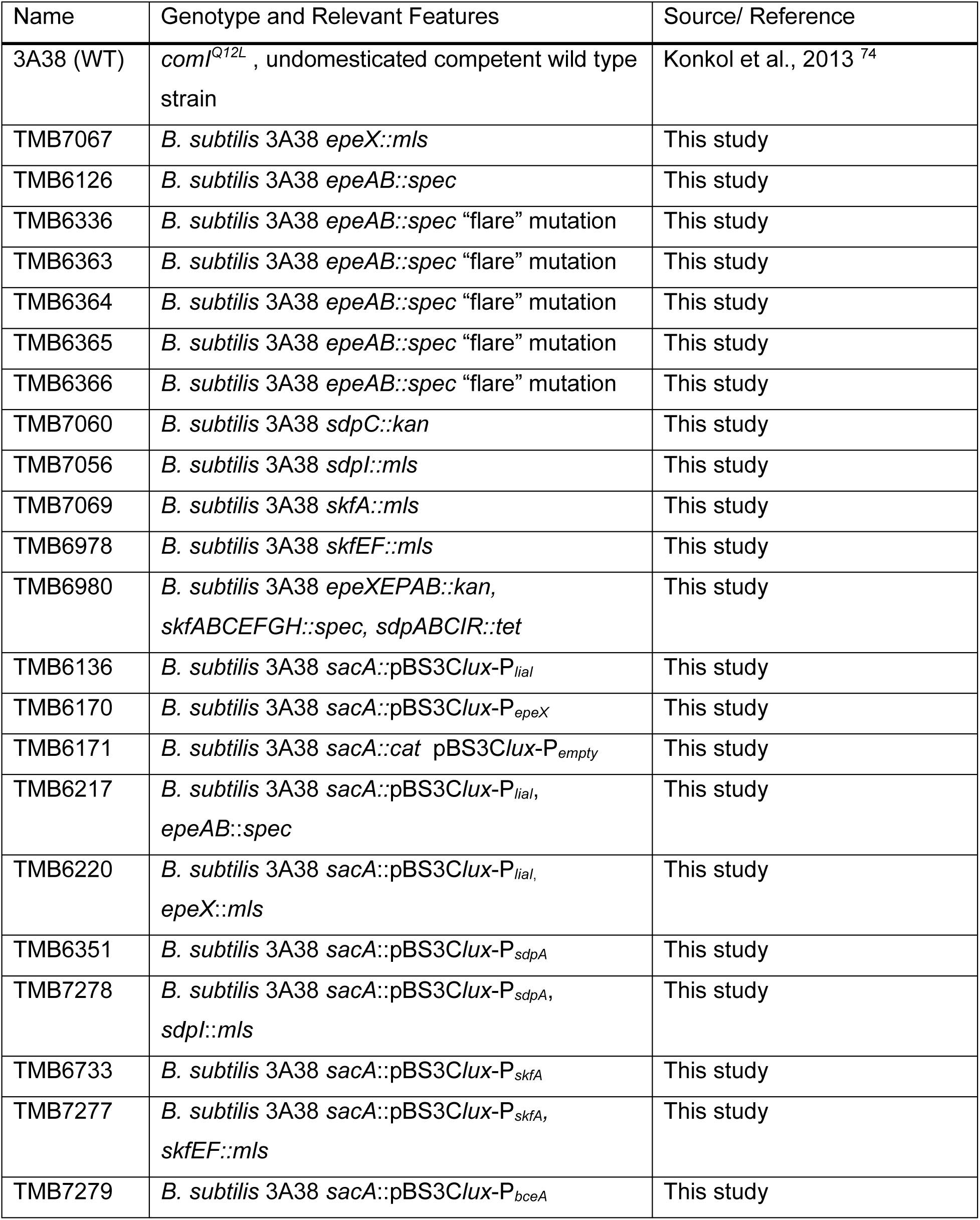
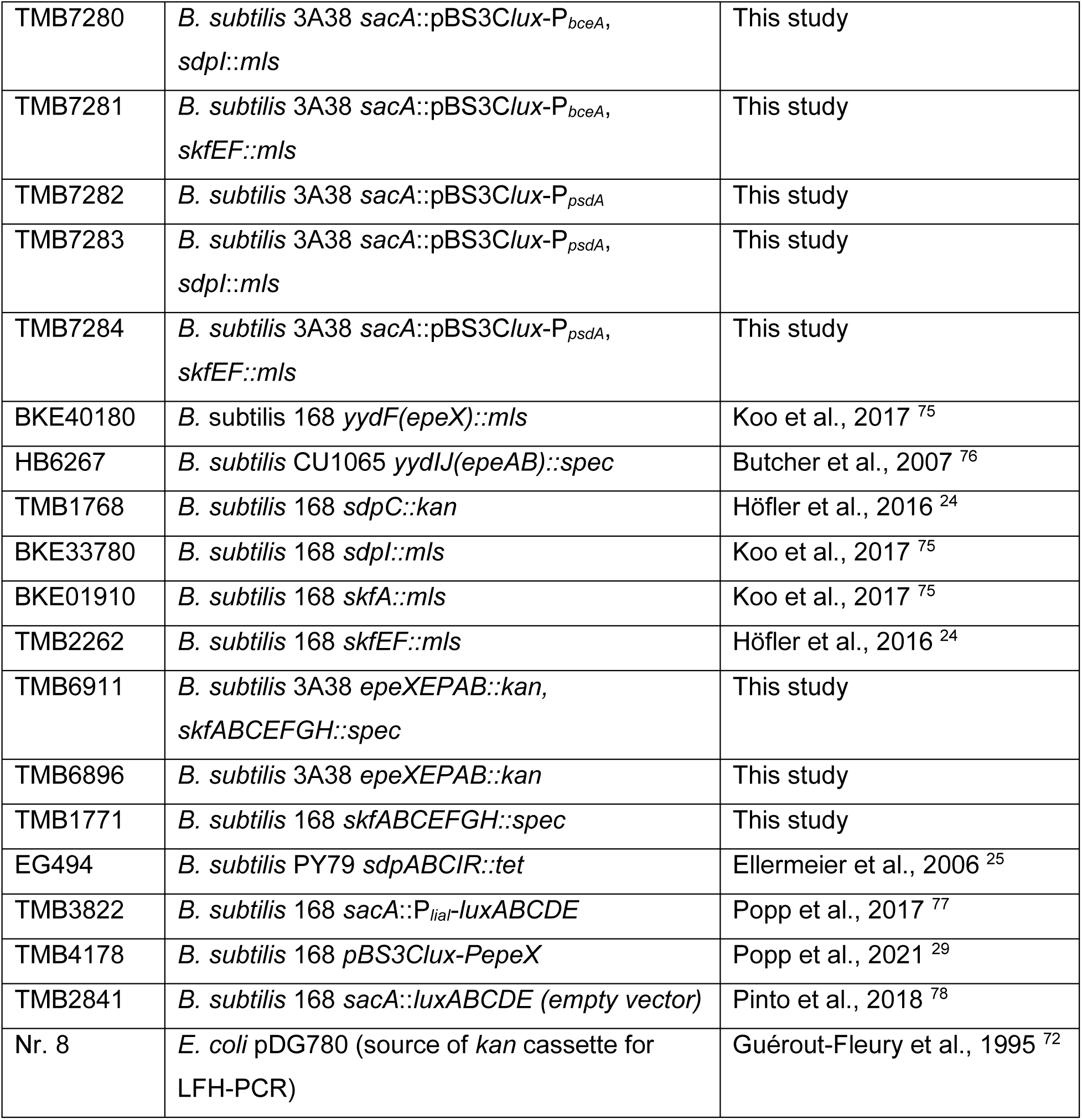
Strains used in this study.

The *epeXEPAB* deletions in *B. subtilis* were constructed by allelic replacement mutagenesis using long flanking homology – PCR (LFH-PCR). The technique is derived from a published procedure ^69^ and was performed as previously described ^70,71^. The kanamycin resistance cassette was amplified from the suitable template vector pDG780 ^72^ using TM0137/TM0138. Two primer pairs were designed to amplify ∼1,000 bp DNA fragments flanking the region to be deleted at its 5′ and 3′ ends (TM7999/TM8000, and TM8001/TM8002). The resulting fragments are here called the “up” and “do” fragments. The 3′ end of the “up” fragment as well as the 5′ end of the “do” fragment extended into the gene(s) to be deleted in a way that all expression signals of genes up- and downstream of the targeted genes remained intact. Extensions of ∼25 nucleotides were added to the 5′ end of the “up”-reverse and the “do”-forward primers that were complementary (opposite strand and inverted sequence) to the 5′ and 3′ ends of the amplified resistance cassette.

All obtained fragments were purified using the PCR purification kit from *Süd-Laborbedarf GmbH* (Gauting, Germany). 100 to 150 ng of the up and do fragments and 250 to 300 ng of the resistance cassette were used together with the specific “up”-forward and “do”-reverse primers at standard concentrations in a second PCR in which the three fragments were joined by the 25-nucleotide overlapping complementary ends and simultaneously amplified by normal primer annealing. The PCR products were directly used to transform *B. subtilis*. Transformants were screened by colony PCR using the up-forward primer with a reverse check primer annealing inside the resistance cassette (TM7999/TM0147). The integrity of the regions flanking the integrated resistance cassette and the cassette itself were verified by sequencing PCR products amplified from chromosomal DNA of the resulting mutants.

All other 3A38 mutants were constructed through re-transformation of the corresponding *B. subtilis* 168 mutants. All source strains were checked by sequencing as described above. Transformation of *B. subtilis* was performed as described previously ^73^, with the addition of 100 µL of chromosomal donor-DNA. Transformants were screened by colony PCR (oligonucleotides, see **Table *1***).

### Growth conditions

The bacterial strains used in the present study are listed in **Table 2**. For routine growth of *B. subtilis*, lysogeny broth (LB) liquid media was made using the following recipe: 1% (w/v) Bacto-peptone, 1% (w/v) NaCl, 0.5% (w/v) yeast extract. For solid plates, LB broth was supplemented with 1.5% (w/v) agar. All LB media were sterilized by autoclaving. For transformation of *B. subtilis*, chemically defined medium MNGE [88.2% 1 × MN medium (1.36% (w/v) dipotassium phosphate × 3 H_2_O, 0.6% (w/v) monopotassium phosphate, 0.1% (w/v) sodium citrate × H_2_O), 1.9% glucose, 0.19% potassium glutamate, 0.001% (w/v) ammonium ferric citrate, 0.005% (w/v) tryptophan, and 0.035% (w/v) magnesium sulfate] was used. Single isolate biofilm assays were conducted using MSgg (Minimal Salts glycerol glutamate) medium. MSgg was made by first preparing a base medium, consisting of 5 mM potassium phosphate, 100 mM MOPS at pH 7.0, and 1.5% (w/v) agar. The base medium was autoclaved and cooled to 55 °C before supplementation with 2 mM MgCl_2_, 700 µM CaCl_2_, 50 µM FeCl_3_, 50 µM MnCl_2_, 1 µM ZnCl_2_, 2 µM thiamine, 0.5% (v/v) glycerol and 0.5% (w/v) glutamic acid. A volume of 25 mL melted MSgg medium was added to each 10 cm square petri dish (*Sarstedt AG & Co. KG*, Nümbrecht, Germany) and solidified at room temperature. The surface of the solid plates was dried for 45 min under a laminar flow cabinet before use in experiments.

*B. subtilis* cells carrying a resistance marker were selected for using spectinomycin (100 μg/mL), tetracycline (12.5 µg/mL), kanamycin (10 μg/mL), or erythromycin (1 μg/mL) combined with lincomycin (25 μg/mL) for MLS.

### Biofilm formation assay

For the cultivation of biofilms, overnight cultures in LB medium with selective antibiotics were diluted to OD_600_ 0.1 in 1 mL LB medium. Carefully, 5 µL of the cell suspension was spotted on pre-dried MSgg agar plates. For MALDI-MS-Imaging analysis and flow cytometry, 5 µL of the cell suspension was spotted onto a 0.22 µm pore size mixed Z-cellulose esters filter membrane (*Merck Millipore*, Darmstadt, Germany) on MSgg agar as previously described ^44^ and incubated at 25 °C. Optical pictures of colony growth were taken daily over the course of 8 days using either a P.CAM360 (*TU Dresden*, Dresden, Germany, see **Fig. S1**) or a binocular microscope WILD M3Z (*Leica Microsystems GmbH*, Wetzlar, Germany, see **Fig. 1**) with a ProgRes “SpeedXT core5” camera (*Jenoptik AG*, Jena, Germany; Capture Pro software). Assays were performed in biological duplicates and technical triplicates.

### Luciferase assay

For luciferase assays on MSgg agar plates, biofilms were grown as previously described herein. Luminescence was detected with a Fusion FX system (*Vilber Lourmat GmbH*, Eberhardzell, Germany). The exposure times for the luminescence reporters were as followed: P*_epeX_* 5 s; P*_sdpA_* and P*_skfA_* 10 s; P*_bceA_*, P*_psdA_*, P*_liaI_* and P*_empty_* 2 min. Assays were performed in biological duplicates and technical triplicates.

### Sample preparation for MALDI-MSI

Biofilms were grown on mixed cellulose ester membranes with 0.22 µm average pore size following the previously described procedure. Membranes were cut into approximately 15–20 mm-wide pieces using a scalpel and removed from the agar surface after 8 days of incubation and subsequently placed in a solution of 10% (v/v) formaldehyde (*Carl Roth GmbH* & Co. KG, Karlsruhe, Germany) in water for 30 min to ensure reproducible full inactivation of microbial viability. Biofilms were then washed two times with ultrapure water and, after drying, mounted on a microscope slide using superglue (*UHU GmbH & Co KG*, Bühl, Germany). A microscopy slide scanner (VS200, *Evident GmbH*, Hamburg, Germany), equipped with an ORCA-Fusion C14440 20UP camera (*Hamamatsu photonics*, Hamamatsu City, Japan) was used for brightfield microscopy of biofilms. For microscope image visualization, OlyVia (version 3.4.1, *Evident,* Hamburg, Germany) was used. A solution of 7 mg/ml 2,5-dihydroxyacetophenone (*Merck KgaA*, Darmstadt, Germany) dissolved in 75% acetonitrile (*Carl Roth GmbH & Co. KG*, Karlsruhe, Germany), 10% methanol (*Carl Roth GmbH & Co. KG*, Karlsruhe, Germany), 10% trifluoroacetic acid (*Carl Roth GmbH & Co. KG*, Karlsruhe, Germany) and 5% ultrapure water was sprayed on the biofilm as MALDI matrix using a SunCollect pneumatic spray robot (*SunChrom*, Friedrichsdorf, Germany). The matrix was applied within 22 spraying cycles at a nitrogen back pressure of 2.5 bar in a meandering pattern. The flow rate followed a gradient from 15 µL/min in the first cycle that increased to 20 µL/min and 30 µL/min for the second and third spraying cycles, respectively. From the fourth spraying cycle onwards, the flow rate was 50 µl/min until the end. The spray nozzle moved with a velocity of 700 mm/min and a line distance of 2 mm. The distance between the spray nozzle and sample surface was set to 44 mm.

All MALDI-MSI measurements were conducted with a timsTOF fleX MALDI-2 QTOF mass spectrometer (*Bruker Daltonics*, Bremen, Germany), equipped with a Smartbeam3D laser emitting at 355 nm. For all MALDI-MS imaging experiments, the resulting field size was 50 μm using beam scan function and the “M5 small” setting with 200 laser shots per pixel with a frequency of 1 kHz. For all MALDI-MSI experiments, a nitrogen pressure in the ion source of 2.5 mbar was used. With the analytical focus on the detection of the peptide toxins, all herein reported data were acquired in the “High sensitivity Detection” mode with the m/z detection range set to 1000 – 4500. All experiments reported here were conducted in positive ion mode and with the MALDI-2-postionization disabled. MSI data was processed using SCiLS Lab MVS (vs. 2024a Pro, Bruker Daltonics, Bremen, Germany) and ion images visualized as false color imaging using the same software. MSI experiments were performed in biological and technical duplicates.

### Microbial flow cytometry and cytometric fingerprinting

Flow cytometric analysis was conducted according to the recently described procedure ^43^. In short: a BD Influx v7 Cell Sorter, (Becton Dickinson, Franklin Lakes, NJ, USA) equipped with a stream-in-air nozzle of 70 µm was used. The blue 488 nm Sapphire OPS laser (400 mW) was used for measurement of the forward scatter (FSC, 488/10; PMT1; related to cell size), side scatter (SSC, trigger signal, 488/10; PMT2; related to cell density), SYTO9 fluorescence (530/40; PMT3), and the PI fluorescence (616/23; PMT5). The UV laser 355 nm Genesis OPS laser (100 mW) was used to measure DAPI fluorescence (460/50; PMT9). Fluidics were run at 33 psi through a 70 μm nozzle and cells were detected equivalent to an event rate of 2,500 to 3,000 events sec−1. The calibration of the cytometer was performed in the linear range by using 1 μm blue fluorescent FluoSpheres (Ref. F-8815, Molecular Probes, Eugene, OR, USA), 2 μm YG fluorescent FluoSpheres (Ref. F-8827, ThermoFisher Scientific, Waltham, MA, USA). For calibration in the logarithmic range, 0.5 μm and 1 µm YG fluorescent FluoSpheres (Ref. F-8813 and F-13081, ThermoFisher Scientific, Waltham, MA, USA) were used. The obtained flow cytometry data were stored in the Zenodo database under the following DOI: https://zenodo.org/records/14967544. Biopsy samples were taken from one biological replicate using a 200 µL pipette tip from six different positions, covering the biofilm from the center to the outer edge. Based on a low cell count, biopsy samples were not OD-adjusted during cell handling. For analysis of population heterogeneity related to cell cycle states, DAPI staining was performed as described elsewhere ^43^. For discrimination between live and compromised cells, a combination of SYTO9 and PI staining was used ^43^. For each bacterial strain, 50.000 cells were measured flow cytometrically and visualized in 2D plots using the program FlowJo 10.0.8.r1 (FlowJo, Becton Dickinson, Franklin Lakes, NJ, USA). Cell gate setting and the creation of the gate template was done as suggested by Abbaszade and colleagues ^43^. FACSorting of different cell types for fluorescence microscopy was performed using the gates from the gate template with 50.000-200.000 cells sorted for each gate. Data analysis was conducted according to previous literature ^43^.

### Fluorescence microscopy

To immobilize spores for microscopy, agarose pads (approx. 20 mm diameter and 1 mm depth) were prepared using 1% UltraPure Agarose (*Life Technologies GmbH*, Darmstadt, Germany) applied to a rubber mold on a microscopy slide and allowed to harden. 2 μL of cell suspension was applied to the agarose pads and covered with a coverslip. The prepared slides were analyzed using a Zeiss AXIO Observer Z1 (*Carl Zeiss AG*, Oberkochen, Germany) with an Axiocam 503 mono (*Carl Zeiss AG*, Oberkochen, Germany) at a total magnitude of 630x (immersion objective PlanApo 63x/ 1.40 Oil Ph3 M27, NA1.4). Microscopy was performed using phase-contrast (Ph3, light intensity: 4.4 V, exposure time: 1.59 s) and BFP-channel (exposure time: 3.13 s for gates: c1n, c2n, c3n, csp1, sp1; 1.81 s for gates sp3, sp4). Excitation wavelength and emission wavelength were 353 nm and 465 nm, respectively. Analysis of the microscopy images was performed using the software Fiji (“ImageJ”). Rectangles were cropped from the original image representative for cells of each cell type.

### Whole genome sequencing

Genomic DNA was isolated with the NucleoSpin® Microbial DNA Kit according to the manufacturer’s protocol. Whole genome sequencing was performed by the Microbial Functional Genomics Group at Ludwig Maximilian University of Munich (Munich, Germany) using Sanger/Illumina 1.9 (paired end read) for analysis of the *epeAB* mutant flare (TMB6336).

Data analysis was performed using the open-source platform Galaxy EU (24.1) (*usegalaxy.eu*, Freiburg Galaxy Team, Freiburg, Germany). For TMB6336, quality of the Illumina paired-end data was verified with “FastQC” (Galaxy v.0.11.8) ^79^. To automate quality and adapter trimming, the sequencing data were analyzed with “Trim Galore!” (Galaxy v.0.4.3.1; threshold: 20; maximum error rate: 0.1; minimum reads length: 20) and sequences bearing an average quality below the threshold value were subsequently removed using “Trimmomatic” (Galaxy v.0.36.5; Average quality required: 20; Number of bases to average across: 4) ^80^. High-quality paired-end data were then assembled as contigs using “Unicycler” (Galaxy v.0.4.8.0; minimum length of contigs: 100 bp) ^81^. Quality check of the assembled genome was performed using “Quast” ^82–84^. Subsequently, the genome assembly contigs were aligned with the genome of *Bacillus subtilis* (ASM904v1) using the “RagTag” tool (reference-guided scaffolding of draft genomes; Galaxy Version 2.1.0+galaxy1) ^85^. For genome annotation, the software tool “Prokka” was used ^86,87^. The assembled genome was aligned to the reference genome to find single nucleotide polymorphisms (SNPs), insertions and deletions using the “snippy” tool with the default parameters (Galaxy Version 3.2) ^88^. For visualization, the script “JBrowse genome” (Version 1.16.11) was used ^89^.

## Acknowledgments

This work was supported by the German Research Foundation (DFG) within the priority program SPP 2389 “Emergent Functions of Bacterial Multicellularity” (Grants K.D. and T.M.: 504017689, S.M.: 503905203). We thank Kim Wüpping (U. Münster) for offering preliminary results of cross sections and enable a more in-depth discussion of the method, Denis Iliasov and Daniela Hartmann for support during WGS analysis using the Galaxy platform, Dagmar Mense and other members of the biomedical mass spectrometry group at U. Münster for help with bacterial culturing and MS data acquisition and evaluation. We are grateful to Bruker Daltonics (Bremen, Germany) for valuable technical support.

## Supplemental data

**Fig. S1.**
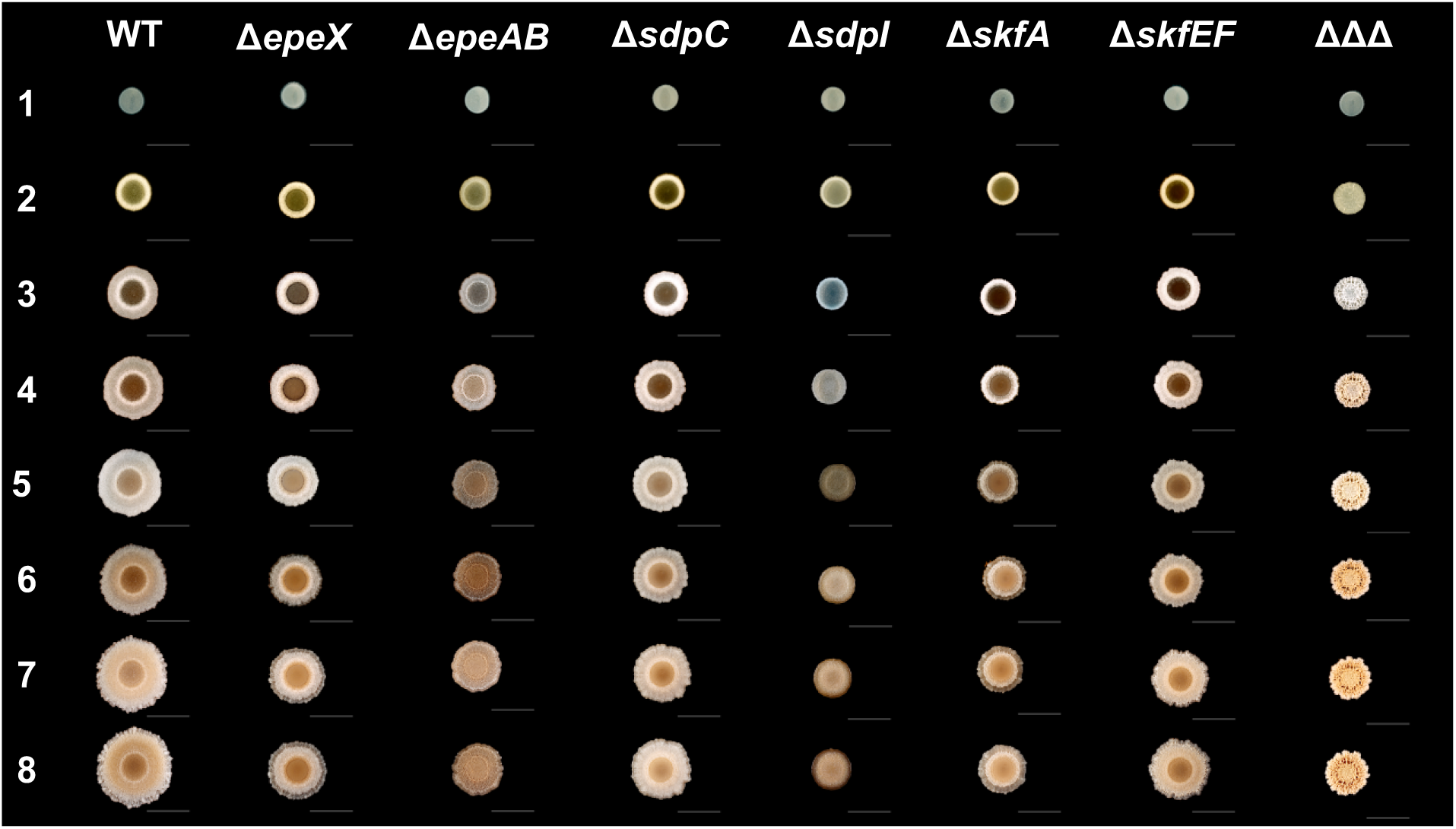
Growth of *B. subtilis* colonies on MSgg agar plates over 8 days. Genotypes of the strains are written above the images, numbers on the left indicate the day of growth. Scale bar indicates 10 mm.

**Fig. S2.**
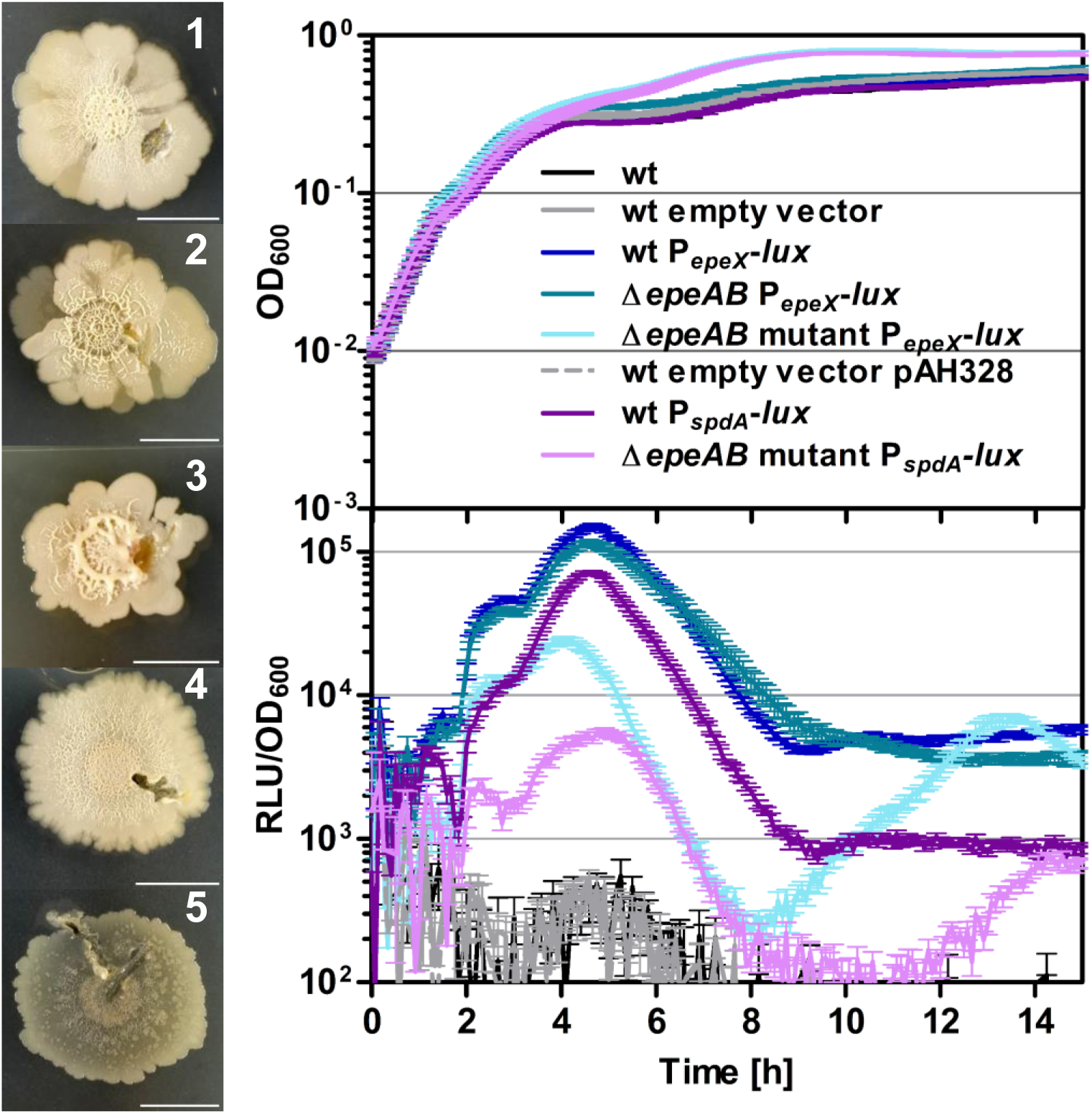
Left: Δ*epeAB* flare mutants 1-5 grown on MSgg agar (day 12 is shown). Right: Growth curve (top) and correlating relative luminescence (bottom) signal over time. Cultures were grown in DSM medium. Color code for each strain is stated within the graph of the growth curve. P*empty*-lux is herein referred to as “empty vector (pAH328)”. Scale bar indicates 10 mm. Error bars indicate standard error of the mean (SEM).

**Fig. S3.**
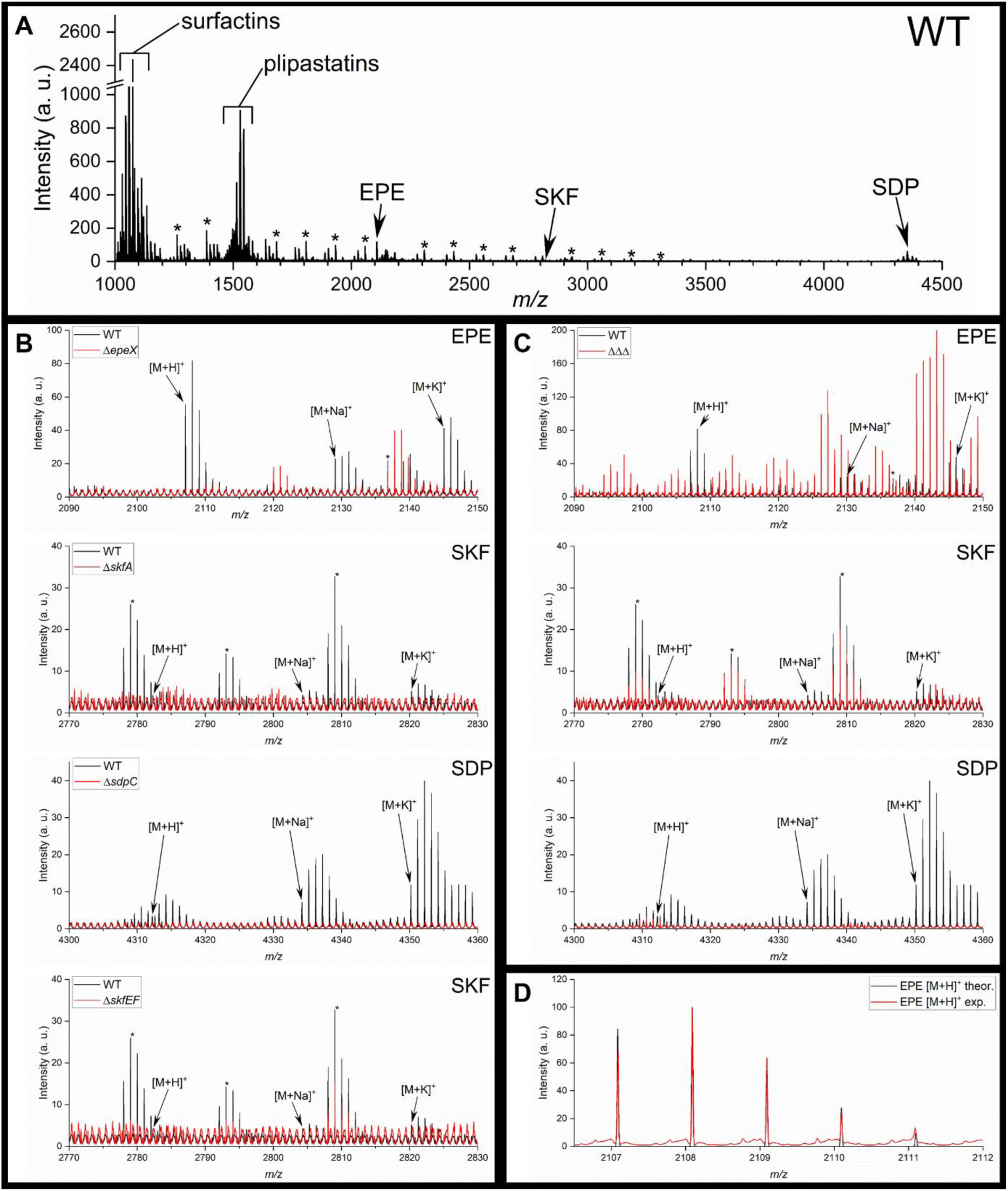
MALDI mass spectra of WT and mutant strains. [A] Overview spectrum of *B. subtilis* 3A38. The signals denoted with an asterisk (*) originate from the adhesive and exhibit a characteristic mass difference of 125.05 Da. This offset corresponds to the molecular mass of ethyl cyanoacrylate, a monomeric component of the adhesive formulation used to fix the colony-coated membranes on the MALDI sample carrier (glass slide). [B] Mass spectra of different single knockout strains (red) in comparison to the WT (black). The lipopeptides are registered as a mix of [M + H]^+^, [M + Na]^+^, and [M + K]^+^ species, as typical for MALDI-MSI in the positive ion mode, and individual molecular species are hence detected as a triplet rather than mono-ionic peak. EPE was recorded at *m/z* values of 2108.09, 2129.08 and 2146.04, respectively; SDP at *m/z* 4314.22, 4334.15, and 4352.18, respectively; and SKF at *m/z* 2783.31, 2804.30, and 2821.27, respectively. [C] Excerpts from the mass spectra of the ΔΔΔ strain showing no cannibalism toxin signals in contrast to WT. [D] Overlay of the calculated isotope pattern of EPE with the experimental values, in this case recorded from a WT colony.

**Fig. S4.**
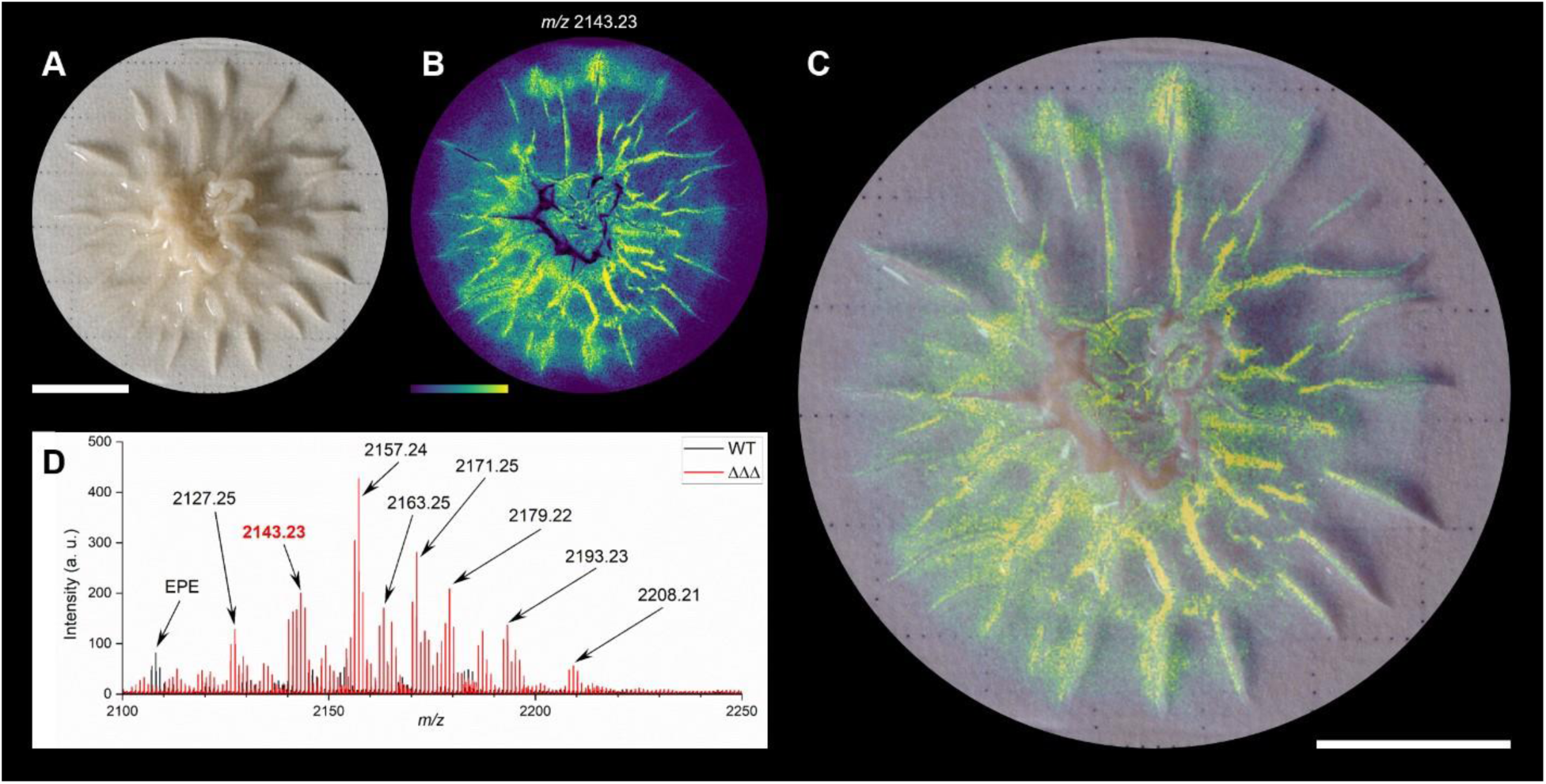
Co-registration of a MALDI-MSI and optical images of a ΔΔΔ biofilm colony. [A] Display of 8 day old ΔΔΔ biofilm grown on the mixed cellulose esters filter membrane. [B] MALDI-MS image of the ion distribution recorded at m/z 2143.23 shown in the viridis color scale; the color scale displays ion intensity from 0% (purple) to 100% (bright yellow). [C] Overlay of both images with 50% transparency of the MALDI image. [D] Zoom-in to the sum mass spectrum recorded from the hypo-cannibalistic biofilm (red trace). Highlighted is the m/z range from 2100 – 2250, displaying a series of hitherto unknowns, most of which are only registered in the mutant. Scale bars indicate 5 mm.

**Fig. S5.**
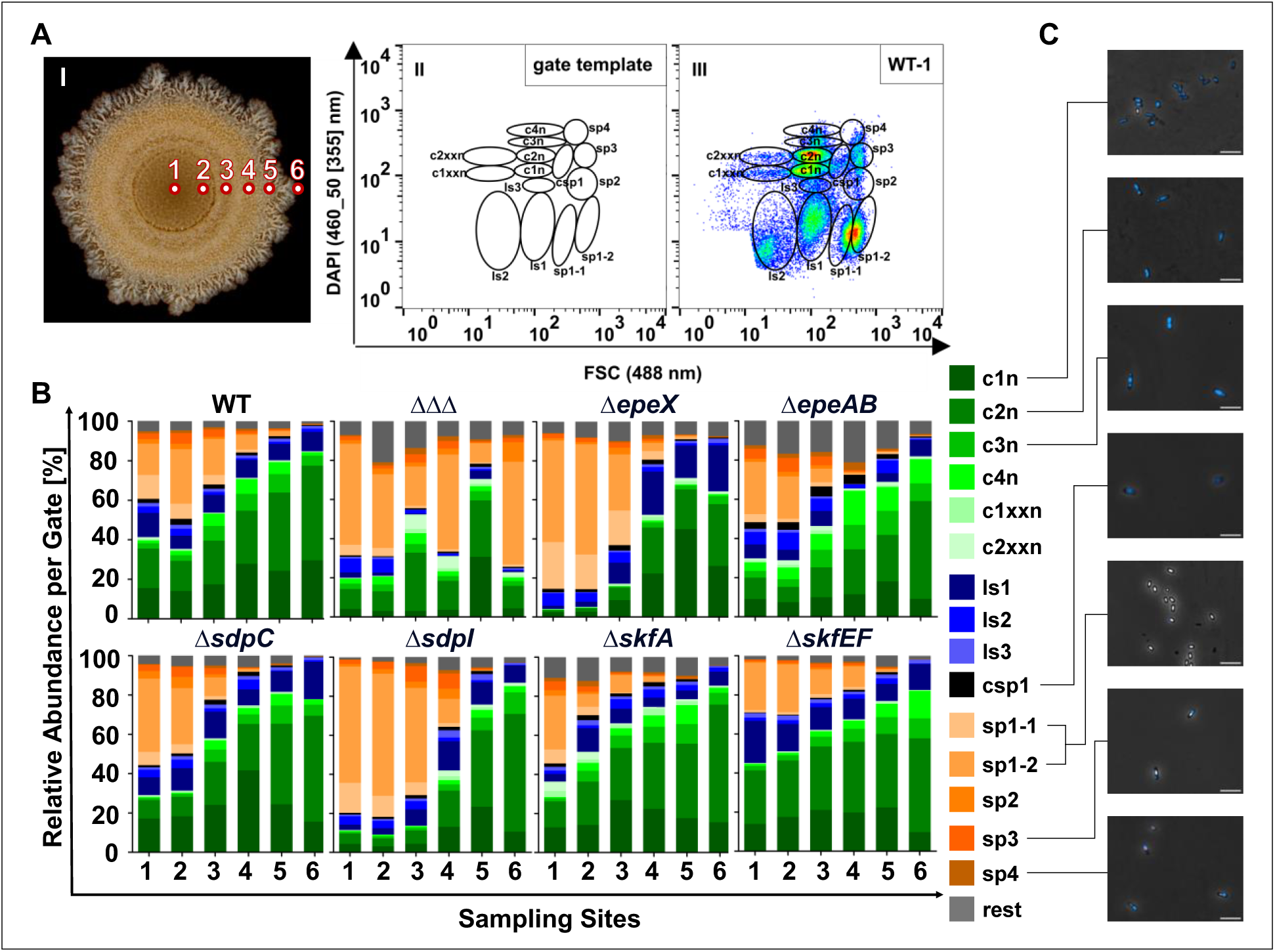
Distribution of cell types across *B. subtilis* colonies regarding cannibalism after DAPI staining. [A] Sampling sites across an exemplary WT colony are shown on the left. On the right, gate templates for analysis and sorting are depicted, as well as an exemplary dataset of flow cytometric measurement of cells present in sampling site 1 of the WT. DAPI signal is plotted against forward scatter (FSC). [B] Overview of flow cytometric analysis results are summarized in bar graphs for each strain, where the relative abundance of cells per gate is plotted against sampling sites. c1n – c2xxn indicate vegetative cells, and ls1 – ls3 indicate less stained cells, meaning that they do not take up DAPI easily. Csp1, sp1-1 – sp4 represent different spore types. [C] Microscopy images of sorted cells from the WT representing some of the analyzed occurring cell types. Scale bar indicates 5 µm.

**Fig. S6.**
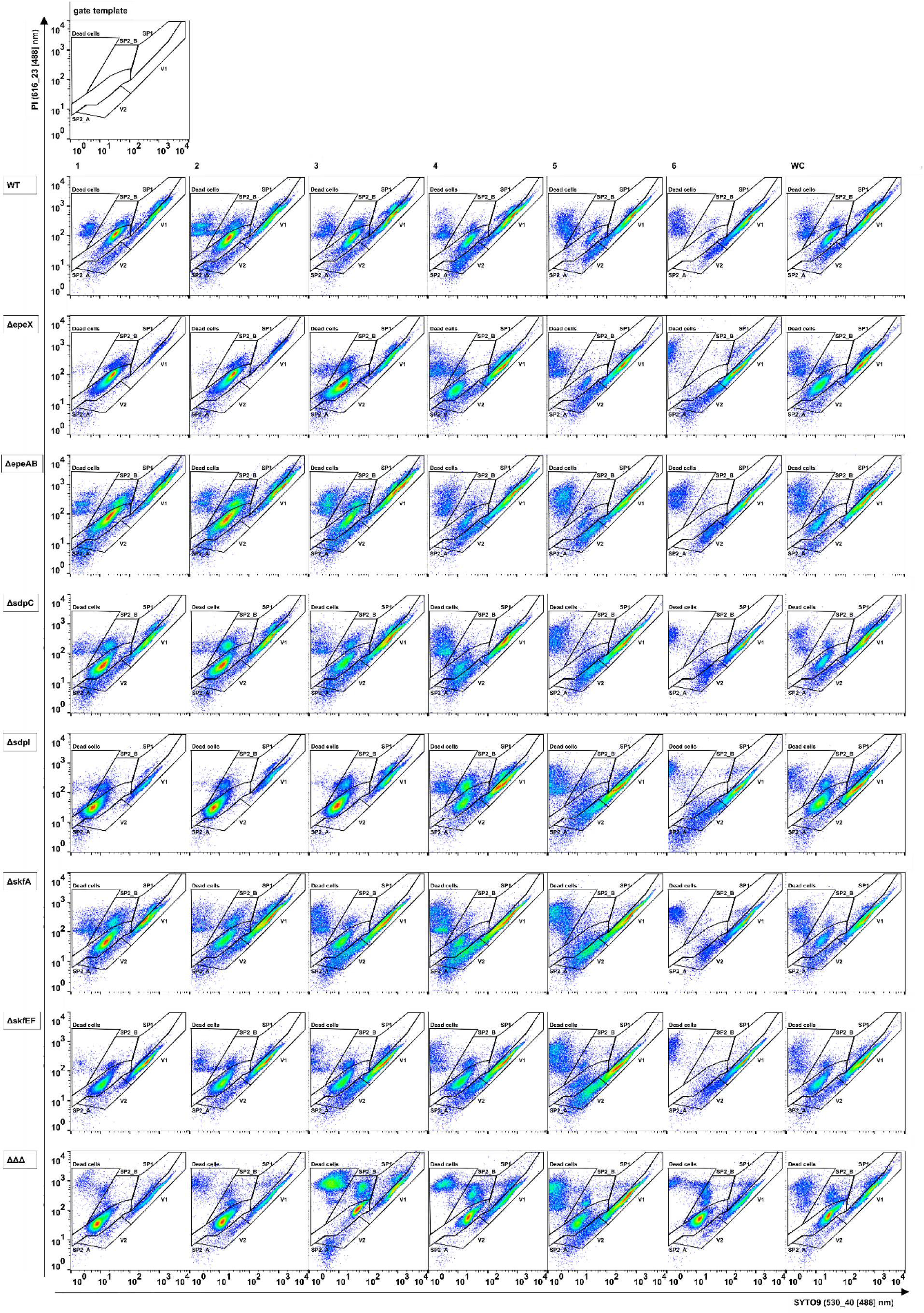
Overview of flow cytometric results for SYTO9 PI staining. Top left shows the gate template. Datasets for each strain are shown in a row, starting with sampling site 1-6, followed by the data obtained for the whole colony (WC). 2D plots depict SYTO9 against PI data [rel. FI].

**Fig. S7.**
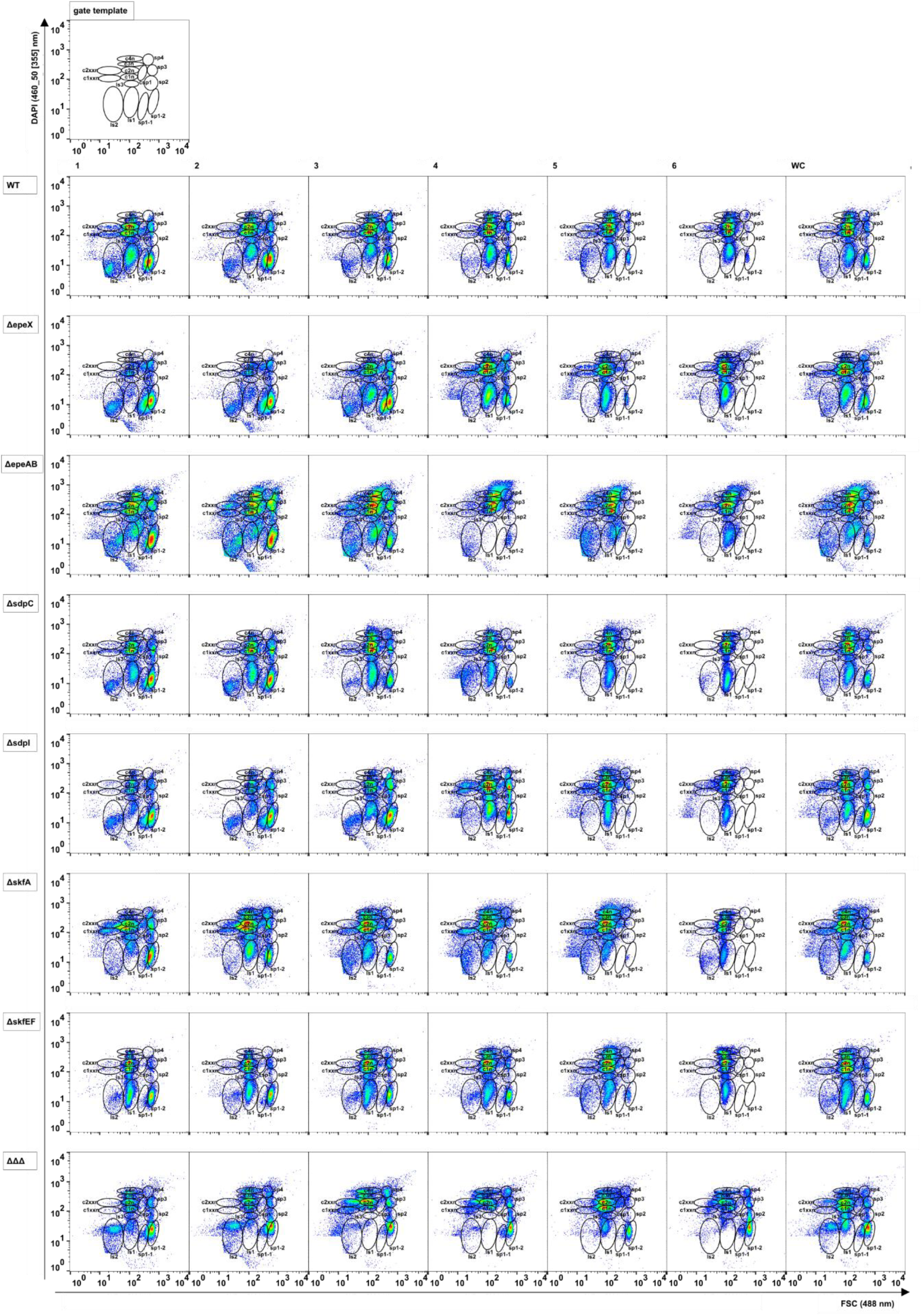
Overview of flow cytometric results for DAPI staining. Top left shows the gate template. Datasets for each strain are shown in a row, starting with sampling site 1-6, followed by the data obtained for the whole colony (WC). 2D plots depict FSC against DAPI data [rel. FI].

